# Trogocytic-molting of T-cell microvilli controls T-cell clonal expansion

**DOI:** 10.1101/2022.05.03.490404

**Authors:** Jeong-Su Park, Jun-Hyeong Kim, Won-Chang Soh, Kyung-Sik Lee, Chang-Hyun Kim, Ik-Joo Chung, Sunjae Lee, Hye-Ran Kim, Chang-Duk Jun

## Abstract

Internalization of the T-cell antigen receptor (TCR) is intimately linked to T-cell activation: a phenomenon thought to be related to the “exhaustion” of T-cell responses. To date, however, no report has considered that during physical interaction with cognate antigen-presenting cells, T cells release many TCRs via T-cell microvilli particles, which are derived from finger-like membrane structures (microvilli) in a combined process of trogocytosis and enzymatic vesiculation and correspond with the loss of membrane TCRs and many external membrane components. Surprisingly, in contrast to TCR internalization, this event leads to rapid upregulation of surface TCRs and remarkable metabolic reprogramming of cholesterol and fatty acids synthesis to meet the demands of clonal expansion, which drives multiple rounds of division and cell survival. We called this event “trogocytic-molting,” which represents an intrinsic molecular basis of T-cell clonal expansion by which T cells gain increased sensitivity to low antigen concentrations.

**TEASER:** “Trogocytic-molting,” led to the rapid upregulation of surface TCRs and tremendous metabolic reprogramming to meet the demands of clonal expansion.

## INTRODUCTION

Many studies over the past three decades have aimed to understand the molecular mechanisms of how the T-cell antigen receptor (TCR) is regulated on the surface of resting and activated T cells, and it has long been thought that TCR downmodulation occurs due to increased TCR internalization, decreased recycling, and increased degradation (*1–6*). However, relatively a few studies have reported that TCR is released from activated T cells in the form of extracellular vesicles (EVs), including exosomes, microvesicles, or lytic granules (*7, 8*). Blanchard and colleagues (*9*) demonstrated that TCR triggering induces exosomes bearing the TCR/CD3/ζ complex in human T cells. Dustin et al. (*10, 11*) demonstrated that T cells produce TCR-enriched microvesicles or synaptic ectosomes at the center of the immunological synapse. Independent of previously published works, Jung et al. and we identified that TCRs are highly enriched in microvilli tips (*12, 13*). Surprisingly, we noticed that microvilli were fragmented into TCR-enriched, nanosized membrane particles (T-cell microvilli particles, TMPs) due to the combined action of two independent mechanisms: trogocytosis and enzymatic vesiculation. Released microvesicles or membrane particles trigger signaling in antigen-bearing B cells (*10*) and dendritic cells (DCs) (*13*), indicating that these nanosized membrane particles can transfer T-cell messages to their cognate APCs. Therefore, we named TMPs “T-cell immunological synaptosomes (TISs)” (*13*).

Interestingly, unlike exosomes, which are secreted from multivesicular bodies, the release of surface microvilli covering the cell outer layer occurs similarly to the manner of molting or shedding, or, in many invertebrates, ecdysis. Molting is the way in which an animal casts off a portion of its outer layer at certain points in its life cycle, typically to allow growth of the organism. In the present study, because T cells also shed their surface microvilli in a combined process of trogocytosis and enzymatic breakage, we termed this phenomenon “trogocytic-molting” or “trogocytic-shedding.” At cellular levels, including in mammalian immune cells, it remains a mystery whether cell growth or division is linked to the trogocytic molting of cellular outer organelles: microvilli.

Although it has long been believed that TCR disappearance on the T-cell surface occurs as the result of internalization and degradation, here we unambiguously demonstrate that trogocytic microvilli shedding is a common process of TCR loss on the T-cell surface during physical interaction with cognate APCs. Surprisingly, we discovered that the components of the TMPs are greatly different from those identified in the T-cell body, and they contain large portions of membrane components, which are essential for TCR signaling and the regulation associated with immune processes. Paradoxically, however, cells that released these essential proteins rapidly recovered surface TCRs and dramatically enforced metabolic rewiring of fatty acids synthesis (FAS), glycolysis, oxidative phosphorylation (OXPHOS), and cholesterol synthesis, leading to multiple divisions and cell survival. By contrast, a condition that internalizes TCRs augmented fatty acids catabolism via β-oxidation (FAO), which could slow or halt the cell cycle progression. Thus, trogocytic molting is an intrinsic regulatory mechanism to control clonal T-cell expansion.

## RESULTS

### T-cell adhesion on cognate APCs triggers TCR internalization and TCR release

*In vitro*, T cells can become activated by several mechanisms, including the use of soluble (sAb) or immobilized (iAb) anti-CD3/CD28 antibodies, causing a significant downregulation of surface TCRs (*14*). To corroborate surface TCR downregulation, CD3^+^ T cells were stimulated with sAb, followed by secondary cross-linking, or plate-immobilized iAb. Surface TCRβ expression was determined at the indicated time points (Fig. 1A), and a dramatic decrease was observed in CD3^+^ T cells stimulated with either sAb or iAb (Fig. 1A).

**Fig. 1.**
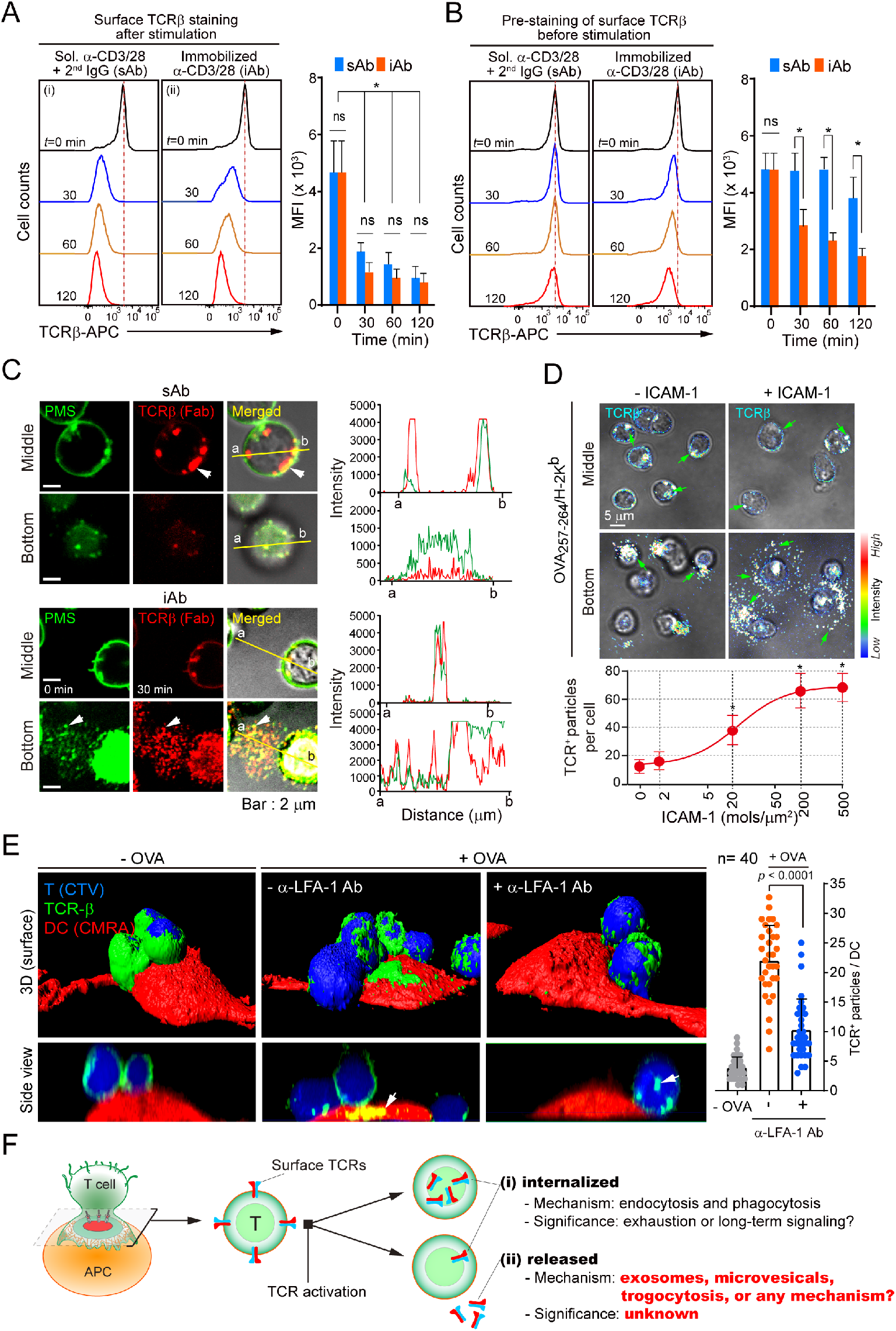
T-cell activation triggers TCR internalization and TCR release. (A and B) Surface TCR expression on primary CD3^+^ T cells. Mouse CD3^+^ T cells were stained with anti-TCRβ (H57Fab-Alexa594) after (A) or before (B) stimulation on plate-immobilized (iAb) or with soluble (sAb) anti-CD3/CD28 antibodies for 3 h. The mean fluorescence intensity (MFI) was measured by flow cytometry. Data represent the mean ± SEM of three independent experiments. ns, not significant (*p* > 0.05), \**p* < 0.05 vs. sAb. C, CD3^+^ T cells stained with the CellMask™ Green Plasma Membrane Stain (PMS) and anti-TCRβ (H57Fab-Alexa594) for 1 h at 4°C, and cells were stimulated as (A) or (B). White arrows indicate internalized or released TCRβ clusters. (D) OTI CD8^+^ T cells were stained as in (B), and cells were placed on a lipid bilayer presenting OVA_257–264_/H-2K^b^/with ICAM-1 (0–500 molecules/μm^2^) for 3 h. Green arrows indicate internalized or released TCRβ^+^ clusters or particles. \**p* < 0.05 vs. 0. (E) OTII CD4^+^ T cells were stained with anti-TCRβ (H57Fab-FITC) and Cell Trace Violet (CTV), and cells were incubated with 1 μg/mL OVA_323–339_-pulsed DCs (CMRA-Orange) in the absence or presence of anti-LFA-1 blocking antibody (10 μg/mL). Arrowheads indicate TCRβ clusters separated from OTII CD4^+^ T cells. TCR^+^ particles per single DC were quantitated using Imaris. Data represent the mean ± SEM of three independent experiments. (F) Schematic model of TCR internalization and release. The model hypothesizes that not all TCRs are internalized (i), but some parts are released (ii) during T–APC interaction.

To monitor whether surface TCRs are endocytosed after T-cell activation, the same T cells were prestained with fluorescence-labeled antigen-binding fragments (Fab), and the cells were stimulated with sAb or iAb (Fig. 1B). Interestingly, TCRβ^+^ fluorescence intensity was significantly reduced in T cells stimulated with iAb, whereas it remained unchanged in T cells treated with sAb (Fig. 1B). The reduction of TCRβ^+^ fluorescence intensity with iAb treatment was not blocked by the endocytosis inhibitor chlorpromazine, the proteasome inhibitor MG132, or the lysosome inhibitor chloroquine (Fig. S1A), suggesting that a significant number of TCRs are released during T-cell adhesion on iAb, although some are internalized.

In fact, T cells become active only after physical contact with cognate antigen-bearing APCs. This suggests that several previously conducted studies overlooked the physiological meaning of TCR release in T-cell biology. Therefore, we monitored the trafficking of TCRβ during T-cell activation. Similar to previous studies (*1, 5, 14–16*), TCRβ^+^ fluorescence signals were dramatically internalized in T cells treated with sAb (Fig. 1C). However, a large portion of TCRβs was released from T cells treated with iAb as particles and spread across the T cells on the plate (Fig. 1C). Interestingly, membrane-specific green plasma membrane stain (PMS) was readily detected in TCRβ^+^ particles, whereas it was internalized in T cells treated with sAb (Fig. 1C, arrowheads). The surface TCR release was also reproduced on the planar lipid bilayers presenting peptide-MHC (p-MHC, ovalbumin [OVA]323–339/I-A^b^), which is specifically recognized by *OTII TCR* CD4^+^ T cells, and recombinant intercellular adhesion molecule (ICAM)-1. In the absence of ICAM-1, TCRβ^+^ fluorescent signals remained on the OTII CD4^+^ T-cell surface with minimal internalization and release. However, large amounts of TCRβ^+^ fluorescent signals were spread out and detected over the plate in a density-dependent manner of ICAM-1 (Fig. 1D). Under physiological settings, DCs also acquired a significant amount of TCRβ^+^ fluorescence signal from OTII T cells, and this event was entirely dependent on the adhesion of T cells (Fig. 1E, white arrowheads, and Video S1–S3). In the presence of anti-LFA-1 Ab, TCRβ^+^ fluorescent particles were not released, but instead internalized into the OTII T cells. To rule out suspicion of anti-TCR Fab artifacts that could be internalized or released in live cells, the similar experiments were performed expressing GFP-tagged TCR-ζ chain (ζ-GFP) in OTII CD4^+^ T blasts (Figs. S1B and S1C). As similar to the anti-TCR Fab, TCR ζ-GFP signals were significantly reduced in OTII CD4^+^ T blasts on iAb but not sAb (Fig. S1B), and released upon binding to the OVA-pulsed DCs (Fig. S1C). Taken together, these results suggest that TCR release is a crucial mechanism to downregulate TCRs at the surface of activated T cells, although some are internalized, as depicted in Figs. 1F (i) and (ii).

### TCR^+^ particles are shed from membrane microvilli during physical contact on immobilized activation matrix

The routes of the release of various EVs are depicted in Fig. 2A. However, exosomes and microvesicles are not likely involved in TCR downregulation because they are released in a contact-independent manner (*17*). The adhesion-dependency of TCR release suggests another third mechanism (*18*). In this regard, we previously reported that T-cell microvilli (finger-like membrane protrusions) are highly fragile membrane structures that can be extracted when two cells split, and are transformed into nano-scale particles (i.e., TMPs) by the actions of a membrane-budding complex, such as TSG101 and Vps4a/b (*13*).

**Fig. 2.**
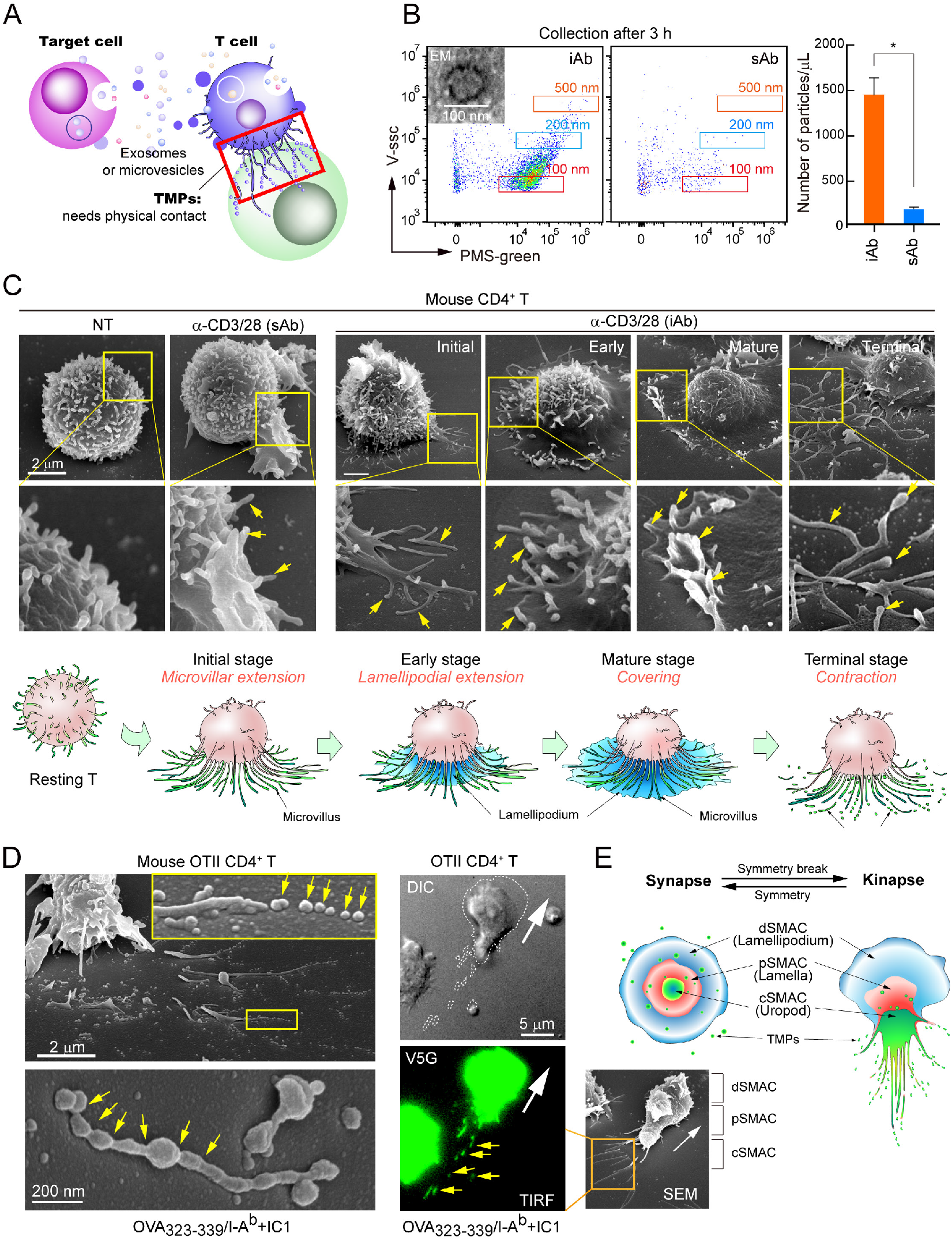
TCRs are transferred to the APC surface during cognate interaction. (A) Schematic diagram describing the generations of TMPs and exosomes or microvesicles. Red box indicates the process by which TMPs are produced. (B) Quantification and size analysis of EVs (produced by iAb or sAb) by flow cytometry. The number of particles per μL was automatically counted using the CytoFLEX Flow Cytometer. \**p* < 0.05 vs. iAb. (C) SEM of naive CD3^+^ T cells stimulated with anti-CD3/CD28 sAb for 3 h or on iAb at various time points. Early (early stage, 1 min), mature (1–5 min), late (10–20 min), and terminal stages (>60 min). A schematic model of TMP generation is presented. (D) Scanning electron microscopy (SEM) evidence of TMP release on the lipid bilayer presenting OVA_323–339_/I- A^b^/ICAM-1 (left). Yellow arrows indicate spherical TISs before and after fragmentation from T-cell microvilli. Release of V5G^+^ microvilli particles from OTII CD4^+^ T cells during migration on OVA_323–339_/I-A^b^/ICAM-1 (right). (E) A schematic model describing TMP generation during T-cell kinapse.

Consistent with the results shown in Fig. 1, quantitative flow cytometry demonstrated that a significant number of particles in the 100–150-nm range were detected in samples generated on iAb-coated surface, whereas very few particles were detected in samples generated from sAb (Fig. 2B). Scanning electron microscopy (SEM) analysis showed no significant changes of microvilli after 3 h stimulation with sAb (Fig. 2C, sAb). However, after initial extension to the iAb surface (Fig. 2C, iAb, ∼ 1 min=initial and early), microvilli remained at the most distal edge of distal-supramolecular cluster (d-SMAC) during maturation of IS (∼5 to 10 min=mature), and then released from T cell body at terminal stage in the form of large rod-shaped TMPs (∼ 60 min=terminal) (Fig. 2C). A schematic model of particle generation is depicted in Fig. 2C. Similar results were also observed in mouse OTII CD4^+^ T cells placed on lipid bilayers presenting OVA_323–339_/I-A^b^/ICAM-1 (Fig. 2D, OTII CD4^+^ T cells in kinapse). Un-fragmented large rod-shaped particles were further fragmented (Fig. 2D, arrowheads). Using the microvilli-specific protein Vstm5 tagged with GFP (V5G, Fig. S2A), we corroborated that long membrane protrusions are fragmented at the rear edge of migrating cells (Figs. S2B and S2C, and Fig. 2D, OTII CD4^+^ T cells in kinapse; Video S4). A schematic model describing particle generation from protrusive rear edges during T-cell kinapse is shown in Fig. 2E.

We obtained further evidence that TCRs are released largely through microvilli-derived particles. First, confocal imaging of TCR ζ-GFP in CD4^+^ T blasts or Jurkat T cells revealed that TCR ζ-GFPs were exclusively localized to the thin microvilli of T cells (Fig. 3A and Fig. S3A). Second, the apical surface of microvilli consists of lipid rafts (*19*), which are sphingomyelin- and cholesterol-enriched membrane signaling domains and are associated with TCR signaling (*19, 20*). In the resting state, TCRs and cholera toxin B subunit (CTB), which clusters ganglioside GM1 and creates lipid rafts (*21*), were significantly overlapped, mostly remaining on the cell surface (Fig. S3B and Video S5), whereas most were internalized under sAb condition (Fig. 3A, Fig. S3B and Video S6). In contrast, both were released and spread around the T cells on the iAb (Fig. 3A, Fig. S3B and Video S7). Third, each microvillus contains F-actin bundles (*18, 22*) and ERMs (ezrin, radixin, and moesin) (*23*). TCRβ^+^ fluorescence signals were co-localized with F-actin and ezrin at resting (NT = no treatment) state but were dissociated from F-actin (Fig. S4) and ezrin (Fig. 3B) through internalization by sAb. However, they were released through the F-actin/ezrin-enriched microvillar protrusions at the late stage of activated T cells placed on iAb (Fig. S4 and Fig. 3B). Fourth, TCR loss correlated with the shedding of microvilli, as determined by SEM (Fig. 3C) and flow cytometry (Fig. 3D). Remarkably, only TCRβ and CD3 complexes were reduced by sAb, whereas several microvilli-specific molecules, including CD62L, GM1, CD4, and CD8, were also reduced by iAb (Fig. 3D) (*23*). A decrease of microvilli-specific molecules was also observed in OTII CD4^+^ T cells on OVA_323–339_/I-A^b^/ICAM-1, whereas they were minimally reduced in the absence of ICAM-1 (Fig. 3D). Fifth, lipid compositions enriched in TMPs and total cell extracts were determined. Based on log fold changes, we identified substantial enrichment of specific lipid species in TMPs, including sphingomyelin (SM) and phosphatidylserine (PS), which were known to be enriched in lipid rafts and exosomes (Kolmogorov-Smirnov tests, p-values < 1 × 10^-5^, Figs. 3E and 3F). Notably, we also found that more enrichment of saturated phospholipids compared to unsaturated counterparts, including phosphatidylglycerol (PG), phosphatidylinositol (PI), and lysophosphatidic acid (LPA), which would increase lipid packing density in TMPs (Wilcoxon tests, p-values < 0.05). Because TMP generation requires both trogocytosis (activation-induced T-cell adhesion) and enzymatic breakage (vesiculation or shedding), we named this event “trogocytic-molting” or “trogocytic-shedding.” Collectively, these results led us to investigate the physiological significance of TCR loss after the trogocytic-molting of microvilli compared with TCR internalization at the initial stage of naive T-cell activation.

**Fig. 3.**
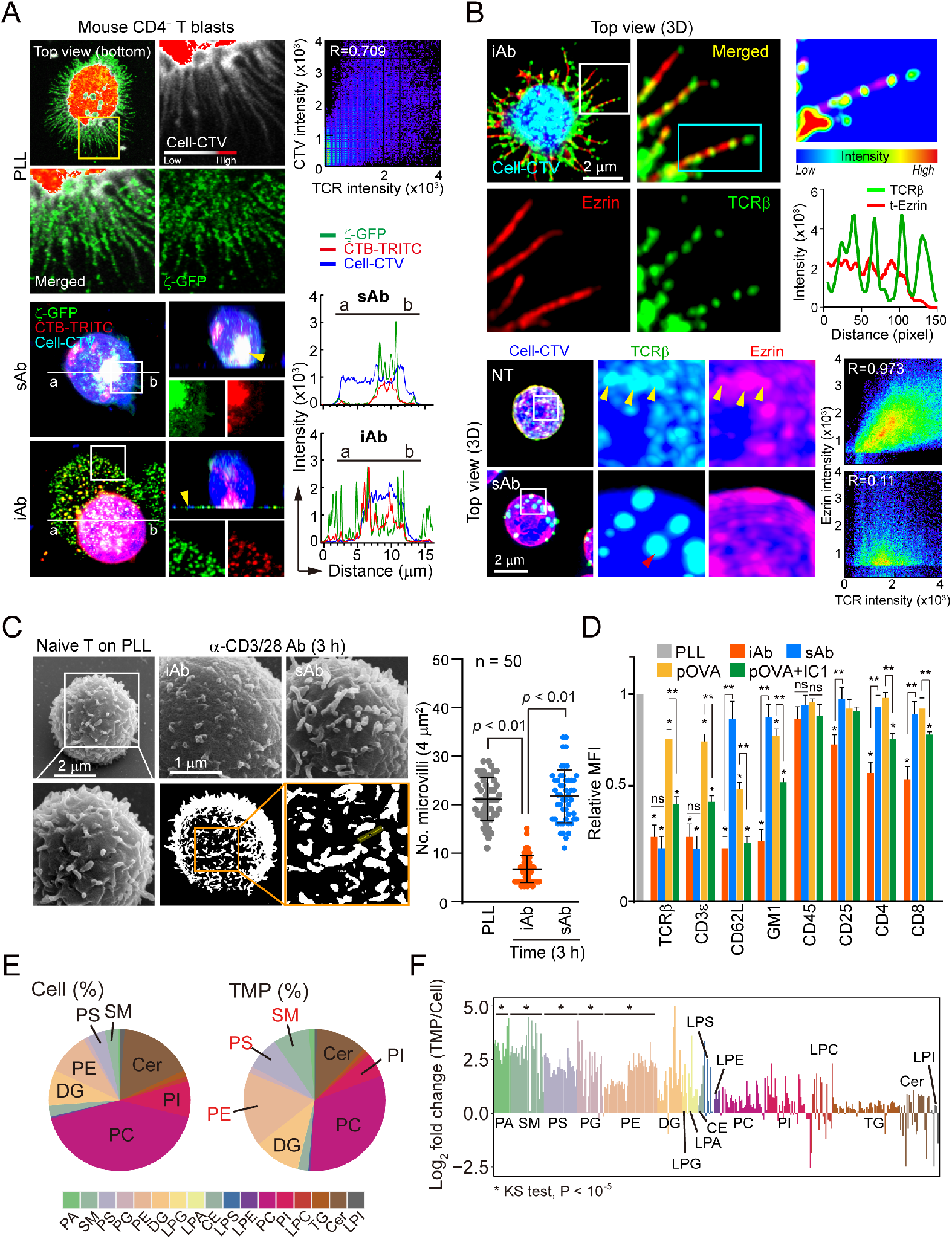
T cells shed surface microvilli when they bind to the adhesion matrix during activation by TCR signaling. (A) TCRζ_GFP expressing CD4^+^ T blasts were stained with CTV or CTB (cholera toxin B) and activated on iAb (10 μg/mL) or sAb (10 μg/mL) for 3 h. Localization of TCRζ_GFP in resting (PLL, top) or activated conditions (TCRζ_GFP or CTB) was examined. Yellow arrows represent internalized or released TCRζ_GFP at each condition. (B) Naive CD3^+^ T cells stained with anti-TCRβ (H57Fab-Alexa594) were stimulated as (A), fixed, permeabilized, and stained with anti-ezrin. Colocalization of TCRβ and ezrin was analyzed by confocal microcopy. (C) Naive CD3^+^ T cells were activated by iAb or sAb (10 μg/mL) for 3 h and subjected to SEM. The number of microvilli per 4 μm^2^ was quantitated using the ImageJ software (n = 50). (D) Flow cytometric analysis of surface proteins potentially enriched or excluded in microvilli after the stimulation of naive CD3^+^ T cells. Data represent the mean ± SEM of three independent experiments. \**p* < 0.05, \*\**p* < 0.001. (E) Pie chart showing the global lipidomes of total cell extracts and TMPs. (F) Lipid compositions enriched in TISs and total cell extracts. Lipids that were significantly enriched in TMPs compared with T cells are denoted by an asterisk (*).

### Molting is associated with TCR expression and metabolic upregulation

Previous reports demonstrated that internalized TCR–CD3 complexes are promptly degraded (*4, 6, 14*). This form of immune regulation may explain the “exhaustion” of T-cell responses that is induced by high antigenic burdens and serves to downregulate immune responses (*2*). From this perspective, in contrast with TCR internalization, we hypothesized that TCR release might increase TCR synthesis to upregulate T-cell responses.

To determine the fate of T cells caused by the differences in TCR internalization and release, we first examined the effects of sAb and iAb because a clear contrast condition could be obtained with this antibody system, i.e., TCR internalization and release. To minimize the issue of TCR signaling intensity, we examined the phosphorylation of downstream signals after the stimulation of TCR under soluble or immobilized conditions and observed that the stimulus intensity was similar (Fig. 4A). In addition, there was no difference in the levels of phosphorylated protein kinase θ (PKCθ), extracellular signal-regulated kinase, and p38 kinase until 3 h of stimulation (Fig. S5A). As TCR release or internalization occurs within 3 h, naive T cells were stimulated for only 3 h (−3–0 h), then washed (0 h), and further cultured for the indicated time periods (0–72 h) with fresh media.

**Fig. 4.**
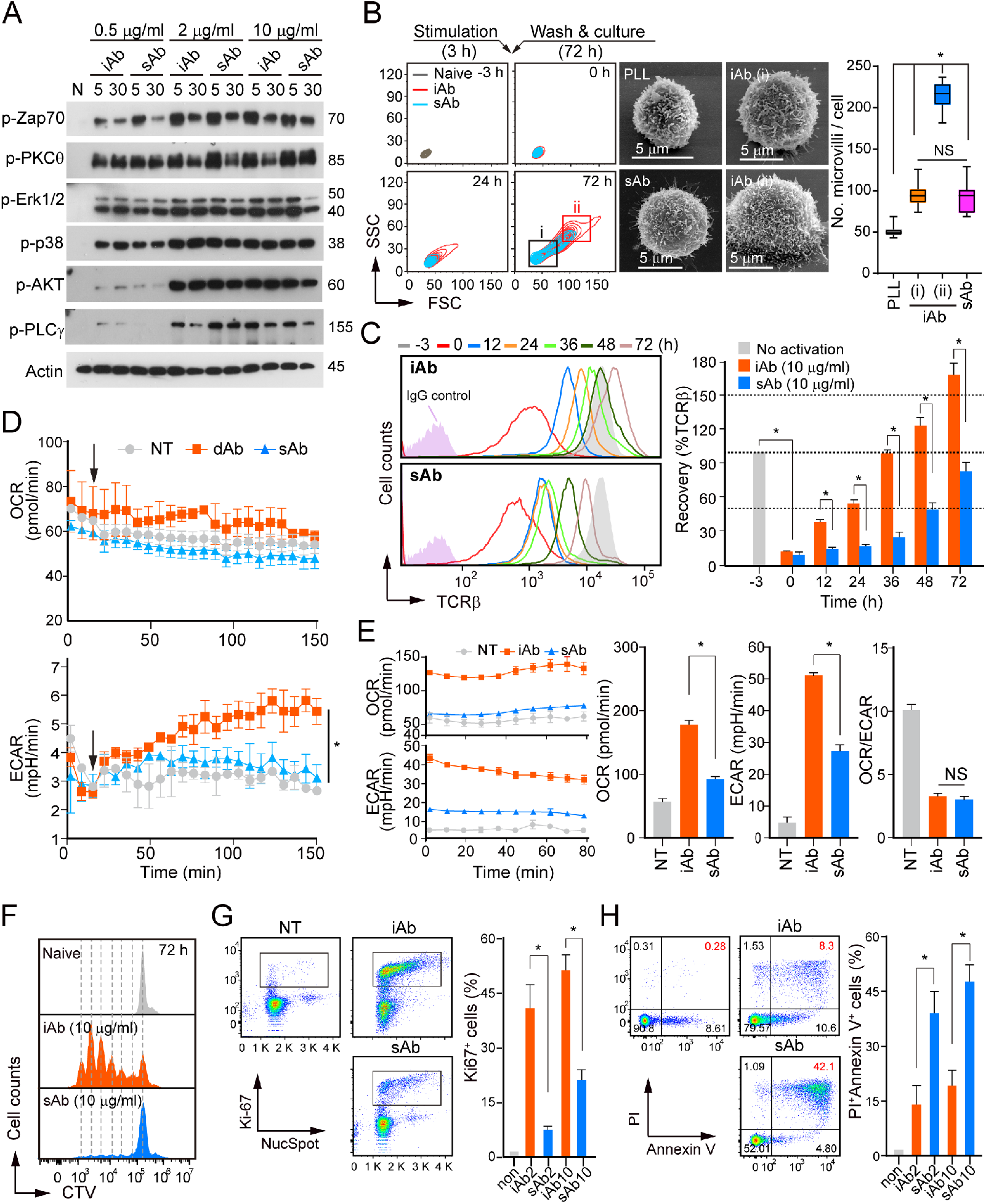
Trogocytic molting of T-cell microvilli enforces metabolic reprogramming for T-cell proliferation and survival. (A) Comparison of the initial TCR signaling strengths between iAb and sAb. (B) Cell size (forward scatter, FSC) or granularity (side scatter, SSC) of naive CD3^+^ T cells stimulated with iAb and sAb. T cells from gating (i) and (ii) were sorted and observed by SEM and flow cytometry. The number of microvilli per cell was quantitated using the ImageJ software. \**p* < 0.05. (C) Surface TCR recovery was determined at the indicated time points during culture. \**p* < 0.05. (D) OCR and ECAR trace of naive CD3^+^ T cells after stimulation with sAb or dAb for 2 h. \**p* < 0.05. (E) OCR and ECAR trace of naive CD3^+^ T cells at 24 h after stimulation with iAb or sAb. Results are representative of three independent experiments. \**p* < 0.05. (F–H) naive CD3^+^ T cells were stained with CTV, stimulated as (B), washed, and further incubated for the indicated time points. Cell division was analyzed at 72 h (F), and proliferation (G) and apoptosis (Annexin V^+^/PI^+^ cells) assays (H) were performed at 48 h after stimulation. NT, none treated; iAb2, 2 μg/mL; iAb10, 10 μg/mL; sAb10. \**p* < 0.05. Data represent mean ± SEM. *Abbreviations*: dAb, dynabead-immobilized antibody; ECAR, extracellular acidification rate; OCR, oxygen consumption rate; PI, propidium iodide.

We initially investigated the shift in the size and granularity of T cells and found a significant difference in their morphology between the two conditions (Fig. 4B). To explore whether TCR expression levels correlated with the number of microvilli and were proportional to the cell size and granularity, T blasts at 72 h of culture were sorted into two groups (i and ii) and analyzed under scanning EM. Group (i) T blasts were similar in the number and size of microvilli as those from sAb (Fig. 4B). However, group (ii) T blasts were larger, contained a greater number of microvilli (Fig. 4B), and exhibited higher expression of surface T-cell proteins than those from group (i) (Fig. S5B). Surprisingly, in contrast to sAb treatment, T cells treated with iAb displayed a rapid recovery of surface TCRβ (Fig. 4C).

T-cell activation results in the metabolic remodeling of naive T cells to a program of aerobic glycolysis, which can generate metabolic intermediates essential for cell growth and proliferation (*24*). We compared the metabolic changes of naive T cells between the two conditions. To mimic the iAb condition, we utilized CD3/CD28-coated dynabeads (dAb). A real-time analysis revealed that the injection of dAb significantly upregulated ECAR, an indicator of aerobic glycolysis, signals (Fig. 4D). A concomitant slight loss of oxygen consumption rate (OCR), an indicator of OXPHOS, was observed under both conditions (Fig. 4D). To determine whether the pattern of changes in ECAR and OCR persists over time after stimulation, ECAR and OCR levels were measured after 24–72 h of culture. We observed higher ECAR and OCR levels in the activated T cells by iAb than by sAb (Fig. 4E), indicating distinguished metabolic reprogramming after TCR loss at the T-cell surface. A dramatic increase in cell division (Fig. 4F) and the proliferation indicator Ki-67 (Fig. 4G) and reduced apoptotic cell death (Fig. 4H) were observed in T cells treated with iAb compared with T cells treated with sAb. No qualitative difference was observed in TCR recovery and proliferation in CD4^+^ and CD8^+^ T cells (Figs. S6A and S6B).

### Molting of membrane microvilli is critical for T-cell clonal expansion

In pursuit of direct evidence that trogocytic microvilli shedding is truly critical for T-cell clonal expansion, we developed several strategies. First, to exclude a potential artifact induced by the plate-immobilized antibody, we used ICAM-1 as an adhesion matrix to induce T-cell adhesion. Unlike sAb alone, the plating of naive T cells on an ICAM-1-coated surface in the presence of sAb significantly induced TCR release (Fig. S7A, green arrowheads). Interestingly, similar to the results under iAb conditions, metabolic reprogramming was dramatically increased (Fig. S7B), followed by rapid induction of surface TCR and increased proliferation of T cells (Figs. S7C and S7D).

Second, because LFA-1, a ligand of ICAM-1, also has a costimulatory function, we used irrelevant anti-CD62L (L-selectin) as a microvilli-specific adhesion matrix. Although TCR^+^ particles were not released on plate-coated anti-CD62L antibody (c-α-CD62L), they were significantly released after stimulation of naive CD3^+^ T cells with sAb (Fig. 5A). Additionally, T-cell metabolism, TCR recovery, and the number of Ki-67^+^ cells were significantly increased (Figs. 5B – 5D).

**Fig. 5.**
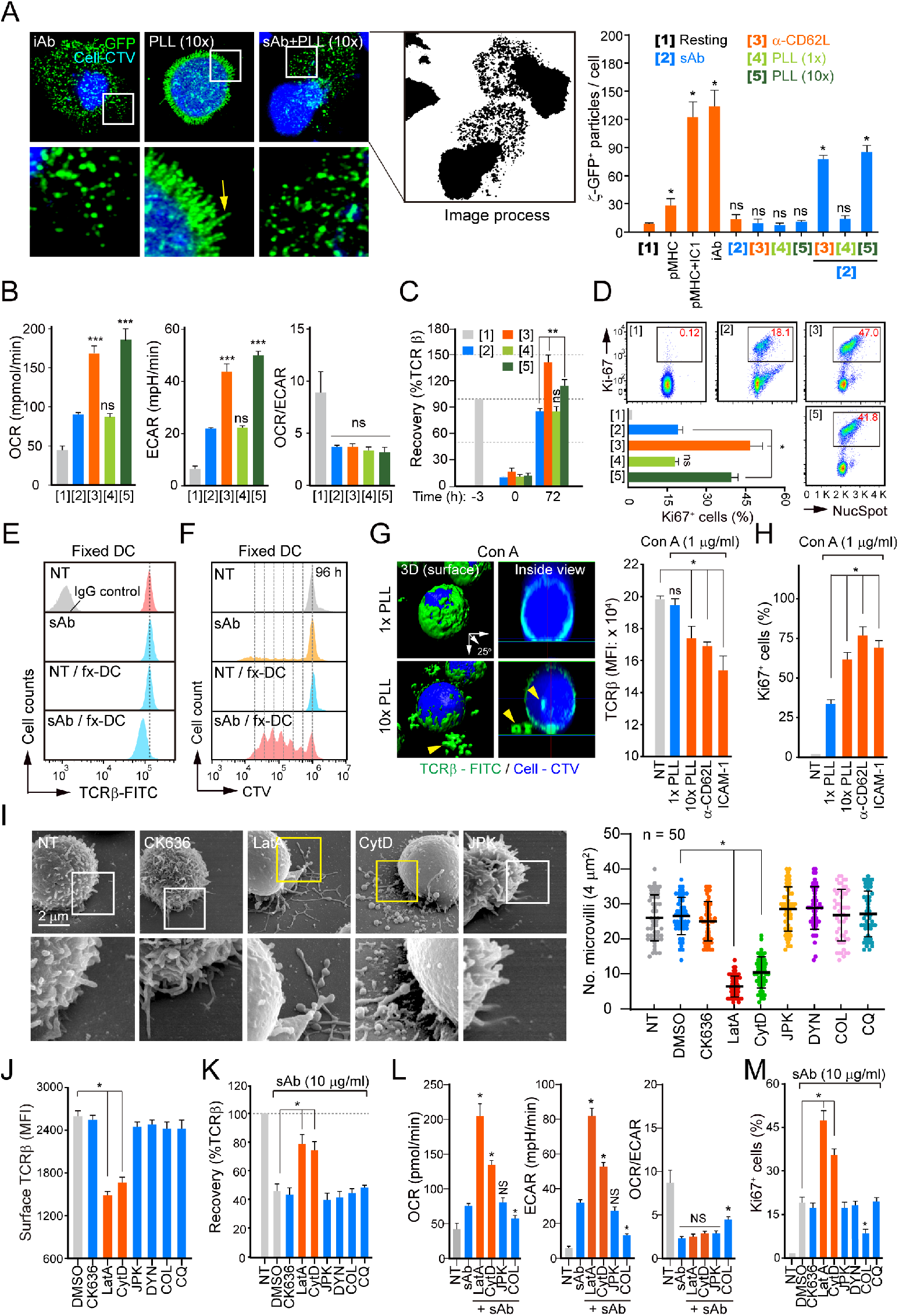
Perturbation of microvilli shedding affects TCR recovery and T-cell proliferation. (A) CTV-labeled CD8^+^ T blasts expressing TCRζ_GFP were stimulated with the indicated stimuli or surface-coated adhesion matrices for 3 h. The distribution of TCRζ_GFP, the number of TCRζ_GFP^+^ particles were examined by Imag J. PLL (1× = 5 × 10^−4^%; 10× = 5 × 10^−3^%), \**p* < 0.05. (B–D), CTV-labeled naive CD3^+^ T cells stained with anti-TCRβ (H57Fab-Alexa594) were stimulated as (A). The OCR and ECAR at 24 h (B), the surface recovery of TCRβ at 72 h (C), and the proliferating populations were determined at 48 h after stimulation (D). Results are representative of three independent experiments. Data represent mean ± SEM. \**p* < 0.05, \*\**p* < 0.01, \*\*\**p* < 0.001. (E–H), CTV-labeled naive CD3^+^ T cells were stained with anti-TCRβ (H57Fab-FITC), incubated with fixed DCs in the presence or absence of anti-CD3/CD28 sAb (E and F), or stimulated with ConA in the presence of indicated conditions for 3 h (G and H). TCRβ intensity on CTV^+^ T-cell surface was measured (E), cells were further incubated for 96 h, and proliferation was analyzed (F). Results are representative of three independent experiments. Distribution of TCRβ_FITC and the MFI of TCRβ (G) were analyzed at 3 h, and proliferating populations were determined at 48 h after stimulation (H). Yellow arrows indicate released or internalized TCRβ. \**p* < 0.05. I – N, Naive CD3^+^ T cells were treated with the indicated inhibitors for 1 h. SEM image, the number of microvilli (I), and TCR levels (J) were analyzed. Cells were activated with sAb in the presence of inhibitors for 3 h, washed, and cultured. TCR recovery at 72 h (K), OCR and ECAR at 24 h (L), and proliferation at 48 h after stimulation (M) were examined. \**p* < 0.05. Values are mean ± SEM.

Third, we investigated whether T-cell activation by sAb also can be increased under even nonspecific adhesion condition. To this end, the plate was coated with a higher concentration (10× = 5 × 10^−3^%) of poly-L-lysine (c-PLL) to immobilize sAb-activated T cells on the culture plate. Strikingly, this condition also increased the release of TCR^+^ particles from sAb-activated T cells, whereas the typical concentration (1× = 5 × 10^−4^%) of PLL showed little effect (Fig. 5A). Again, this condition significantly enhanced T-cell metabolism and resulted in rapid TCR recovery and a greater number of Ki-67^+^ cells (Figs. 5B – 5D). In another experiment, DCs were lightly fixed with 0.4% paraformaldehyde to provide only adhesion sites for T cells. This condition also mimicked the matrix-coated surface (Figs. 5E and 5F).

Fourth, we examined other T-cell stimuli such as concanavalin A (ConA) and phorbol 12-myristate 13-acetate (PMA)/ionomycin instead of antibodies. ConA is known to irreversibly bind to glycoproteins on the cell surface and commit T cells to proliferation (*25*). Interestingly, ConA alone had little effect on TCR internalization, whereas it significantly induced the release of TCRβ^+^ TMPs on the coated matrices (Fig. 5G). Consistently, T-cell proliferation was enhanced (Fig. 5H). Similar results were also observed with PMA/ionomycin (Fig. S8).

Fifth, if the reason for the increase in T-cell metabolism depends on microvilli shedding rather than adhesion itself, the reagents that induce microvilli release or disruption may enhance the T-cell metabolic programming under sAb conditions. We used various inhibitors that interfere with the actions of the cytoskeleton or endocytosis. Among the various inhibitors, latrunculin A (LatA) and cytochalasin D (CytD), both actin-depolymerizing reagents, significantly reduced the number of microvilli and artificially released a significant number of small microvilli particles on the plate (Fig. 5I). In contrast, CK636 (an Arp2/3 complex inhibitor), jasplakinolide (JPK; an actin-stabilizing agent), and other inhibitors had little effect (Fig. 5I). The shedding of microvilli correlated with the reduced TCR levels on the T-cell surface (Fig. 5J). To our surprise, T cells cultured for 48 h after activation with sAb in the presence of LatA or CytD rapidly recovered the T-cell surface TCRs (Fig. 5K) and exhibited enforced metabolic reprogramming for T-cell proliferation (Figs. 5L and 5M), suggesting that surface microvilli release is linked with metabolic reprogramming for T-cell clonal expansion. However, the lack of effect of various endocytosis process blockers suggests that endocytosis of TCR does not correspond to T-cell proliferation.

Sixth, long, rod-shaped microvilli particles are further vesiculated by the actions of a budding complex, such as Vps4a/b (*13*). In fact, TMP contains higher amounts of Vps4a/b (Fig. 6A). We investigated whether knockdown of Vps4a/b may retard the TCR recovery and T-cell proliferation. Si-RNA targeting Vps4a/b significantly reduced the production of TMPs on OVA/H2K^b^/ICAM-1 (Fig. 6A). After stimulation on the lipid bilayer (OVA/H2K^b^/ICAM-1), although TCRβ loss on the surface of OTI T cells was less than that of the scrambled si-RNA-transfected T cells, TCRβ recovery was significantly delayed (Fig. 6B). Similarly, attenuated metabolic reprogramming, followed by a reduced Ki-67^+^ cell population, was observed in T cells (Fig. 6C). Interestingly, CD25 and CD69 expression was not altered by si-Vps4a/b (Fig. 6D), indicating that the reduction of Vps4a/b has no effect on initial T-cell activation.

**Fig. 6.**
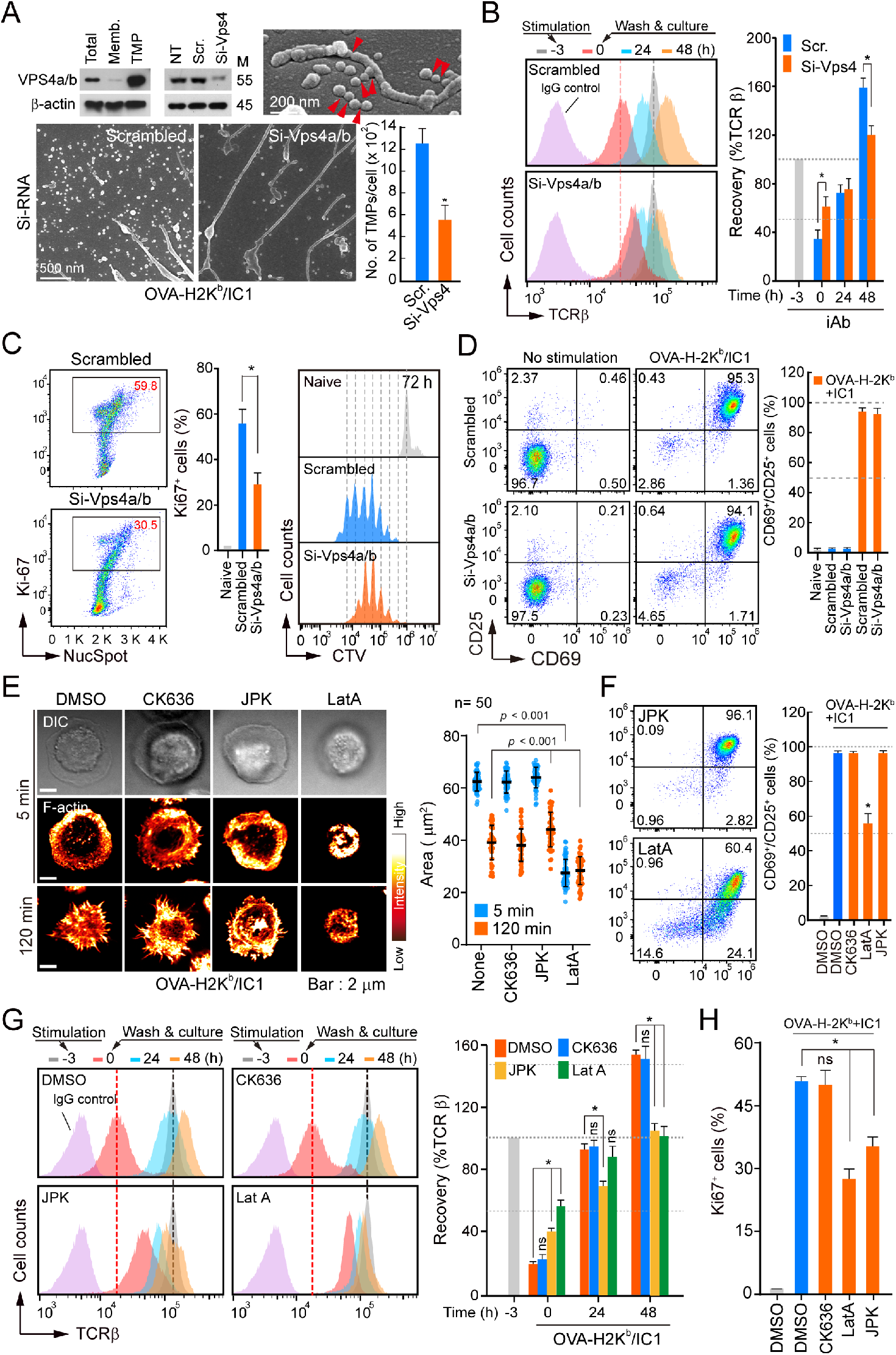
Inhibition of TMP release reduces T-cell metabolism and proliferation. (A–D) OTI CD8^+^ T cells were transfected with si-RNA targeting Vps4a/b for 24 h, stimulated on a lipid bilayer presenting OVA_257-265_/H2K^b^+ICAM-1 for 3 h, and subjected to western blotting or SEM (A), TCR recovery (B), Ki-67 and CTV intensities for proliferation (C), and CD25/CD69 for activation (D). Results are representative of three independent experiments. \**p* < 0.05. (E–H) Naive CD3^+^ T cells were stimulated on a lipid bilayer presenting OVA_257– 265_/H2K^b^+ICAM-1 in the presence of CK636 (100 μM), JPK (100 nM), or Lat A (237 nM) for the indicated time, fixed, and stained with phalloidin-TRITC. Cell spreading was observed by confocal microscopy, and the area was measured using the ImageJ software (E). T-cell activation at 24 h (F), TCR recovery (G), and proliferating populations were determined at 30 h after stimulation (H). \**p* < 0.05.

If actin disruption induces microvilli shedding, then actin stabilization may inhibit microvilli fragmentation, thereby blocking TCR loss. To this end, we tested whether several actin drugs used in this study could affect T-cell activation. CK636 and JPK had little effect on T-cell spreading and subsequent activation on the OVA-H2K^b^/ICAM-1 (Figs. 6E and 6F). Interestingly, the actin stabilizer JPK inhibited the surface TCR loss and resulted in retarded TCR recovery and cell proliferation (Figs. 6G and 6H), but it did not alter the downstream signals of TCR until 3 h of stimulation (Fig. S5A). Notably, although LatA and CytD enhanced T-cell proliferation under the sAb condition (Fig. 5M), they significantly reduced T-cell spreading and subsequent T-cell activation on the OVA-H2K^b^/ICAM-1 (Figs. 6E and 6F). These results suggest that intact F-actin structure, i.e., microvilli composed of F-actin, is critical for T-cell activation during IS formation. However, after proper TCR stimulation, a molting process is essential for T cell proliferation.

### TMPs exclusively contain external membrane components involved in immune and metabolic processes

To elucidate a potential mechanism for how trogocytic molting of microvilli enhances T-cell proliferation, purified TMPs obtained from activated naive CD3^+^ T cells (3 h on iAb) were subjected to liquid chromatography-tandem mass spectrometry (LC-MS/MS). Proteins that were commonly detected from three independent experiments on the total cell lysates (TCLs) and TMPs were selected (Fig. 7A). This analysis revealed 3,537 (80.7%) proteins in the TCLs and 3,216 (74.4%) in TMPs, of which 2,375 (54.2%) were common in both samples. The high degree of overlapping proteins observed between TCLs and TMPs indicated that TMPs contain most of the TCL proteins (Fig. 7A). However, a cellular component analysis based on Gene Ontology annotation revealed that TMP-only exclusively contains external membrane components while TCL-only includes a variety of internal cellular components (Fig. 7B). In addition, analysis and visualization of biological processes with Cytoscape demonstrated that proteins included in TMP-only are related to the regulation of immune processes and vesicle transport while those in TCL-only play a variety of roles in cellular pathways (Fig. 7C).

**Fig. 7.**
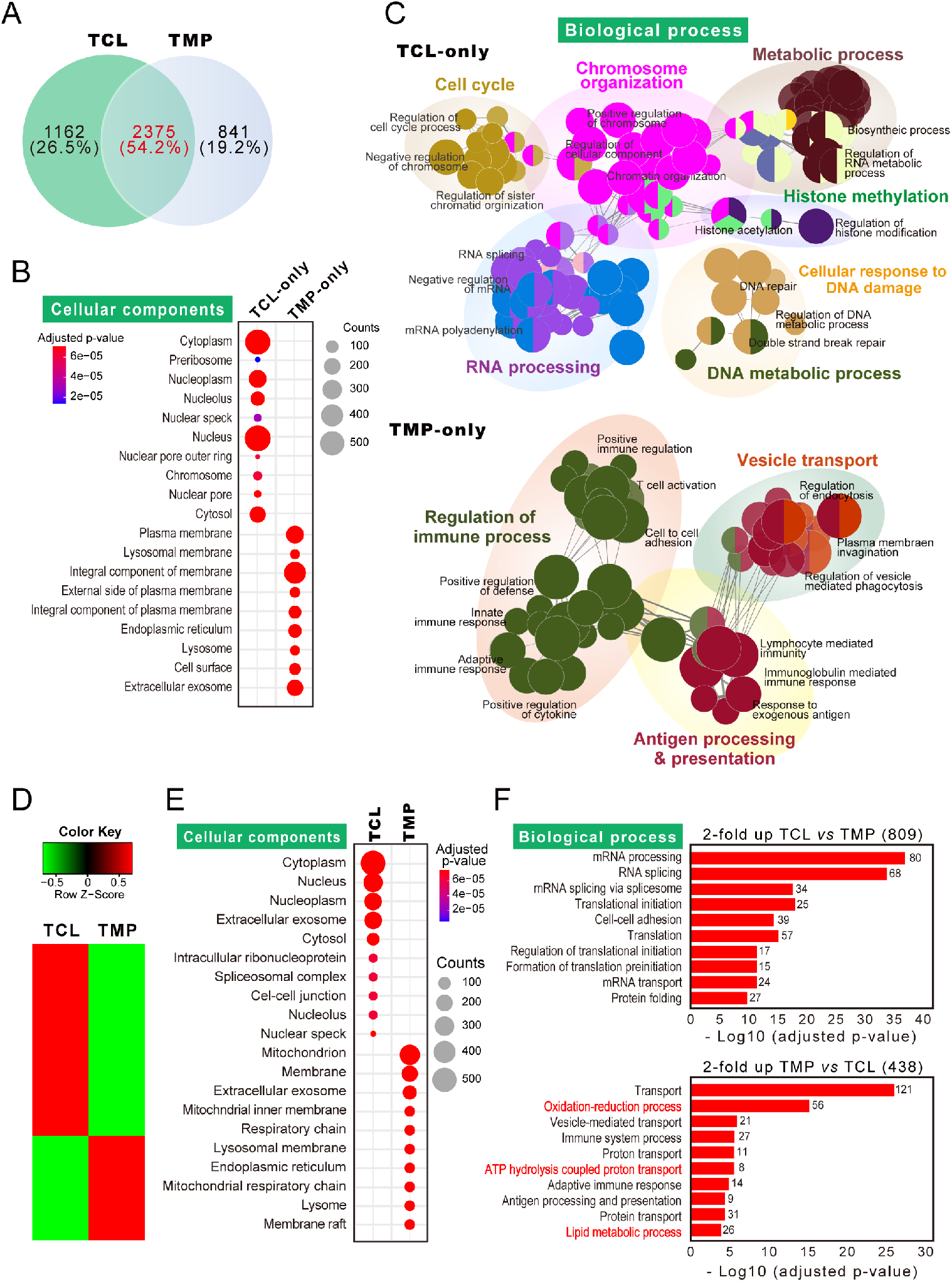
TMPs contain external membrane components involved in immune and metabolic processes. (A) Venn diagram showing the overlap of proteins between TCL and TMP. (B) Analysis of the GO cellular components for the proteins exclusively identified in TMP (TMP-only) or TCL (TCL-only) using DAVID bioinformatics resources 6.8 (https://david.ncifcrf.gov/). (C) Overrepresented GO network for the proteins of TMP-only and TCL-only using Cytoscape V3.6.1 (http://cytoscape.org/) with the ClueGO V2.5.7 plugin. Each node represents a GO biological process, and the size of the node reflects the enrichment significance. Node color represents the class that they belong, and mixed color indicates multiple classes. Only significant interactions (*p* < 0.05) and grouped terms are shown. The two-sided hypergeometric test yielded the enrichment for GO, and Bonferroni step-down correction for multiple testing controlled the p values. (D) Heatmap showing the relative expression pattern of 1247 differentially expressed proteins in TCL and TMP. Proteins included in these analyses met a cutoff of |fold change| >2. (E and F) Analysis of GO cellular component and GO biological process for the proteins in (D) using DAVID bioinformatics resources 6.8. Benjamini–Hochberg adjustment was used for multiple test correction.

We further analyzed the cellular components and biological processes of the proteins commonly found in both groups (2,375 proteins). To this end, particular repertories of proteins were selectively sorted into TCL or TMP based on 2-fold difference by relative quantitation provided by the exponentially modified protein abundance index (emPAI) (Fig. 7D). Similar to the results of TMP-only, cellular component analysis revealed that proteins enriched in TMPs are derived from membrane components (Fig. 7E). Interestingly, analysis of the biological process revealed that, in addition to protein involvement in the regulation of immune function, TMPs contained proteins that are related to energy metabolism, such as ATP hydrolysis, oxidation-reduction process, and lipid metabolic process (Fig. 7F). From the analysis of proteins in TMPs, we further hypothesized that the release of external membrane components such as TCR complex and many T-cell proteins, cholesterols and glycolipids, and some metabolic proteins via shedding of T-cell microvilli may evoke a significant alternation in the process of T-cell activation, and hence T-cell fate.

### T-cell molting favors FAS, whereas TCR internalization favors FAO

To understand whether the release of membrane components, including TCRs, via the molting of microvilli regulates gene expression involved in certain pathways, we generated the transcriptome of naive T cells stimulated with sAb, iAb, LatA, and sAb plus LatA (sAb+LatA) through expression profiling of the microarray (Affymetrix Mouse Gene ST 2.0). Based on hierarchical clustering, we observed the coordinated expression changes by time (baseline, 3 h, and 24 h) and group (naive T, LatA, sAb, sAb+LatA, and iAb-activated T cells) (Fig. 8A). Based on principal component analysis (PCA), we observed the trajectory of expression changes from naive T to LatA, sAb, sAb+LatA, and iAb T cells (red arrow) (Fig. 8B). Focusing on naive T cells compared with sAb- and iAb-treated T cells (Fig. 8C), we performed pathway enrichment tests (R gProfileR package) between iAb and sAb (both 3 and 24 h) (Fig. 8D). Interestingly, we found that the cellular metabolic process was significantly changed at 3 h, whereas the cell cycle process was significantly changed at 24 h (enrichment test p values <10^−8^). Therefore, we speculated that iAb-activated T cells could undergo dramatic metabolic reprogramming at 3 h, and later its reprogramming affected the cellular proliferation at 24 h.

**Fig. 8.**
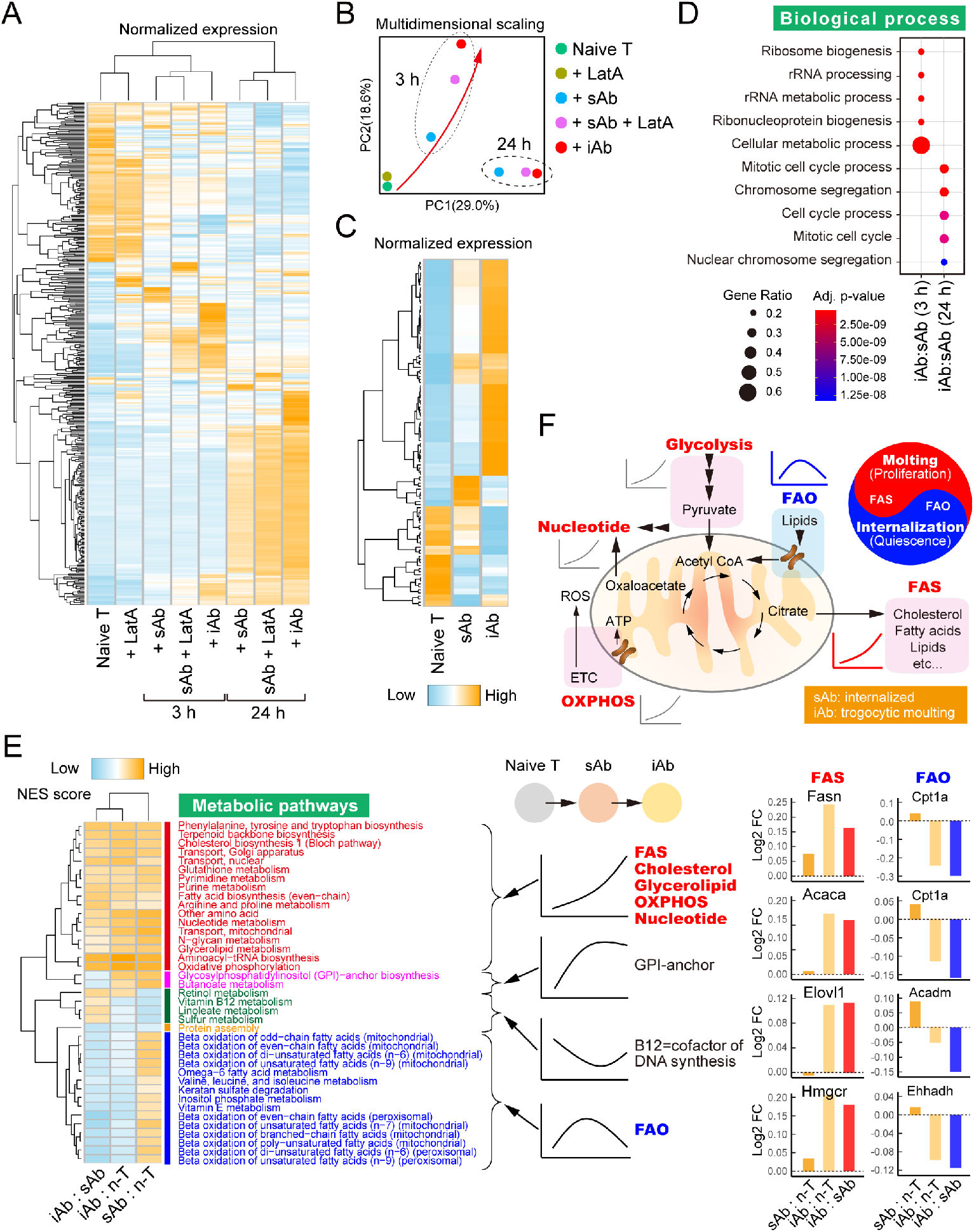
Transcriptome analysis of naive T and sAb-, sAb+LatA-, and iAb-activated naive T cells. (A) Coordinated changes in gene expressions (Affymetrix Mouse Gene ST 2.0 microarray) from naive T (0 h) and LatA-, sAb-, sAb+LatA-, and iAb-activated T cells at 3 and 24 h. Genes on the heatmap were selected based on fold changes from naive T cells (>1.5) and plotted with normalized gene expressions (z-score). (B) Principal component analysis (PCA) of the transcriptome. Trajectory was changed from naive T cells to sAb-, sAb+LatA-, and iAb-activated T cells. The changes were time-dependent, observing different clusters by time. (C) Selected heatmap of naive T and sAb- and iAb-activated cells at 3 h. The group of genes behaved differently, mostly reversed between naive and iAb-activated cells. Genes on the heatmap were selected as described in (A). (D) Enriched biological pathways (Gene Ontology) of differentially expressed genes. (E) Enrichment tests of metabolic pathways of T cells. NES of metabolic pathways (R fgsea package) were calculated from the fold changes of group comparisons. Pathways of the heatmap were selected based on enrichment test p values (< 0.05). Bar graphs show the fold changes (log2) of representative genes in the FAS or FAO pathways. *Abbreviations*: Fatty acid synthesis, FAS; fatty acid oxidation, FAO; Glycosylphosphatidylinositol, GPI; and oxidative phosphorylation, OXPHOS.

The dramatic changes in the cellular metabolic process (Fig. 8D) led us to perform in-depth metabolic pathway enrichment tests using high-quality metabolic maps (*26, 27*). Based on normalized enrichment scores (NES) of expression changes (R fgsea package), we detected distinct clusters of metabolic pathways enriched in differently by conditions. First, we observed the pathways of maximal expressions in iAb-activated T cells, including FAS, OXPHOS, glycerolipid, cholesterol, and nucleotide metabolism, which could promote the recovery of membrane loss and lipid rafts (Fig. 8E). GPI-anchor synthesis, which is also involved in the formation of lipid rafts, was highly expressed in both sAb- and iAb-activated T cells compared with naive T cells (Fig. 8E). Cholesterols and fatty acids are a vital component of cell membranes, can supply substrates for cell signaling, and provide a high-yielding energy source (*24, 28*). Significant upregulation of lipid metabolism pathways (Fig. 8E), together with the increased glycolysis and OXPHOS pathways (Figs. 4D and 4E), in iAb-vs. sAb-activated T cells strongly suggest that membrane microvilli molting is evidently linked with metabolic processes for T-cell proliferation. Second, we detected the following two distinct pathway clusters specifically changing in sAb-activated T cells: 1) a decreased cluster that includes B12 metabolism, a cofactor of DNA synthesis, and 2) an increased cluster that includes FAO, a metabolic feature of memory T cells. Vitamin B12 deficiency reduces proliferation in some cell types (*24, 28*). Therefore, increased FAO and reduced B12 pathways in sAb-activated T cells suggest that the TCR internalization event is linked to slowing or halting of cell cycle progression. The representative genes clustered in the FAS or FAO pathways are shown in Fig. 8E. Interestingly, increased expression of metabolic pathways was also observed in sAb+LatA-activated T cells compared with sAb treatment (Figs. S9A and S9B).

We also generated the transcriptome of naive T cells and ConA (low-dose)-, PLL (10×)-, and ConA-PLL-treated T cells and found that only ConA+PLL, which dramatically release TCR^+^ TMPs, induced the upregulation of cell proliferation (enrichment test p values <10^−8^) (Figs. S10A and S10B) and cellular biomass synthesis (e.g., cholesterol, nucleotide, amino acids, and fatty acids) (Figs. S10C and S10D; fGSEA p values <0.05). Interestingly, genes associated with FAO were not significantly regulated with ConA treatment, presumably because ConA did not cause internalization of the TCR complex (Fig. 5H).

Collectively, the present study demonstrates that the physical interaction of T cells with their cognate APCs is physiologically significant in two manners. First, as shown in previous studies, T-cell activation induces TCR internalization, which can induce “biomass conversion into energy,” thereby slowing or halting T-cell proliferation. Second, at the initial stage of naive T-cell activation, however, T cells require massive expansion to procure effector functions, and this is possibly obtained by the molting of TCR-enriched microvilli, which can induce “biomass generation” (FAs, cholesterols, glycerolipids, etc.). A schematic model based on global metabolic gene profiling and the metabolic functional changes caused by molting or TCR internalization are depicted in Fig. 8F.

## DISCUSSION

TCR downmodulation after engagement with p-MHC is a hallmark of T-cell activation and is believed to occur through increased TCR internalization or rapid degradation (*14, 29*). However, in the present study, we showed that molting of T-cell membrane protrusions including microvilli in the form of TISs is a crucial mechanism corresponding with the loss of TCRs on the surface of activated T cells. T-cell microvilli have at least two major functions. First, they are important outer-layer organelles to sense antigen peptides on APCs (*12, 13, 18, 30*). Second, during the physical interaction with APCs, T-cell microvilli act as carriers to release a large number of proteins and lipids required for T-cell life. Paradoxically, these T cells enter multiple division cycles and grow to meet the demands of clonal expansion by the *de novo* synthesis of gene clusters involved in lipid metabolic pathways, including FAS, cholesterols, glycerolipids, and lipid rafts. To our knowledge, these findings are new evidence that, apart from external factors such as cytokines released from APCs, T cells have an intrinsic program for proliferation and clonal expansion by which the T cells gain increased sensitivity to low concentrations of antigen.

Unlike the many studies focusing on TCR internalization, none has investigated whether the release of TCRs from T cells encountering their cognate APCs can affect surface TCR downregulation. Therefore, no one has examined what percentages of TCRs are internalized or released during T-cell contact with an APC. Interestingly, most TCR-triggering studies that have used surface-coated anti-CD3 antibody or peptide-pulsed APCs did not account for microvesicles or TMPs released from activated T cells. This presupposes that all TCRs are internalized during T-cell activation; thus, the results produced by this preconception are likely to be interpreted from only one perspective. The present study reveals that TCR traveling is distinguished based on whether the stimulating conditions occur in a soluble or matrix-coated state. From this point of view, the coated surface is more similar to the physiological state. Indeed, we (*13*) and others (*31*) have clearly demonstrated that a considerable number of TCRs are transferred to the APCs during IS formation and deformation.

What are the mechanisms of TCR release from T cells encountering their cognate APCs? Unlike the generation of exosomes or microvesicles, microvilli shedding depends on trogocytosis (*17, 32, 33*), which was thought to transfer membrane patches containing intact molecules via enzyme-independent cleavage (*33*). However, we recently identified that fragmentation (or vesiculation) of microvilli from the T-cell body requires enzyme complexes such as TSG101 and Vps4a/b, which mediate the budding of microvesicles at the cell surface (*34*). In an agreement with this line, knockdown of Vps4a/b significantly reduces TMP release via microvilli shedding (*13*). To our knowledge, our work showed the first evidence that T cells release microvilli particles through the combined action of two independent mechanisms: trogocytosis and enzymatic breakage. Therefore, the “trogocytic molting” of surface membrane components may be a common mechanism occurring during immune cell-to-cell interaction.

Interestingly, there has been no attempt to measure the size or volume of trogocytosed materials. To exclude any potential controversy, we compared the quantity of the particles (or exosomes) produced from activated T cells under two different stimuli conditions: iAb or sAb. Compared with the small number of vesicles produced after sAb stimulation, the generation of large number of 100–150-nm small particles from T cells after iAb stimulation within relatively short period (∼3 h) strongly suggests that TCR^+^ particles are not likely released from multivesicular bodies. More clearly, previous reports demonstrated that lipid rafts, which are membrane signaling domains associated with TCR signaling (*19, 20*), are located at the apical surfaces of microvilli (*19*). Interestingly, the apparent presence of CTB, a marker for GM1 in lipid rafts, with plate-bound TCR^+^ particles released from activated T cells on iAb strongly demonstrates that TCR-enriched TMPs are formed from the shedding of surface microvilli. Unlike the sAb condition, a reduced number of microvilli and downregulation of microvilli-specific proteins in T cells activated via the iAb condition also provide a direct evidence that the particle origins are microvilli covering the outer layer of the cell.

Recently, Dustin’s group reported that T cells release TCR-enriched microvesicles or synaptic ectosomes through IS while interacting with APCs (*10, 11*). They demonstrated that membrane particles are primarily released from the central region of the supramolecular activation cluster (c-SMAC). If TCR^+^ particles are solely released at the c-SMAC, the particles must be concentrated in the central region of IS. However, T-cell membrane particles were spread around the T-cell periphery on the matrix-coated surface. Moreover, TCRs were clearly localized through the microvilli stems and tips, which are enriched with ezrin and F-actin, and were released through long rod-shaped fragments. Furthermore, it is now generally accepted that microvilli play an important role in sensing, particularly at the interface with the APCs (*12, 30, 35*).

Of great interest is that most studies related to exosomes or microvesicles have focused only on the effect of these small vesicles on the target cells, tissues, or microenvironments, but none to our knowledge has yet explored whether the release of EVs has an autocrine regulatory function. From this perspective, it is striking that TISs contain, in addition to TCRs, a large number of membrane microvilli or lipid raft proteins essential for T-cell life. Hence, there are two questions, viz., is the molting process unique to T cells, or do other cells also undergo molting during their lifespan? and if molting occurs, what biological or pathological phenomenon is it related to? Scientists in cancer research have believed for several years that the secretion of EVs could be one of the primary mechanisms by which cancer stem cells interact with other tumor and non-tumor cells (*36*). However, no one has evaluated whether EV release is a self-regulatory mechanism by which cancer cells permanently grow. In this regard, it is interesting to note that interfering with EV biogenesis and/or release inhibits cancer cell survival and growth (*36*). Furthermore, extensive evidence suggest that EVs are involved in the metabolic switch occurring in cancer and tumor stroma cells. If molting is a general phenomenon for cell proliferation, it may also be important in stem cell research. There is evidence that stem cells use plasma membrane protrusions as platforms for EV shedding (*37*).

In general, organisms will become exhausted and eventually die if they lose components essential for sustaining their life cycle. However, a natural phenomenon for which such a paradox exists is in molting, shedding, or ecdysis, in which a creature casts off a part of its body to grow. In this perspective, an important question is how membrane shedding was molecularly linked to enhance metabolic reprogramming for cell proliferation and survival. Since microvilli are enriched with membrane essential lipids, such as cholesterols, glycerolipids, and FAs, which are necessary for T-cell metabolism and subsequent activation and proliferation, the loss of such essential lipids via TMPs is likely to promote the *de novo* synthesis of gene clusters for lipid metabolism. This pathway may be linked to trigger Notch1 signaling, which is known to be involved in T cell proliferation and survival (*38*). Additionally, no energy burden in using ubiquitin machinery to clear unnecessary receptors or signaling molecules may also be linked to Notch1 activation, thereby inducing the downstream cell cycle regulator c-Myc (*38*). In addition to membrane components, TMPs contain significant amounts of mitochondrial proteins. The lack of energy-related mitochondrial proteins may promote the synthesis of genes related to the OXPHOS pathway. Overall, the present results of LC-MS/MS-based lipidomics and proteomics and transcriptome analysis strongly suggest that T cells release molecules that are essential for antigen recognition (TCR and its complex), energy metabolism (glycolysis, TCA cycle, and ATP binding proteins), and membrane components (FAs and cholesterols), which paradoxically promote T-cell proliferation for clonal expansion.

Approximately 20 years earlier, the Schoenberger group reported that naive cytotoxic T cells require a single short interaction (within ∼2 h) with stimulatory APCs to initiate a program for autonomous clonal expansion and differentiation into functional effectors (*39*). However, the mechanism of why a brief period was sufficient remained an enigma for a long time. In the present study, we have compiled extensive evidence that short-term shedding of T-cell membrane protrusions is a critical mechanism for initiating a program of T-cell clonal expansion. Importantly, an increased number of microvilli and increased TCR expression are markers for antigen-experienced or memory T cells. Further research is underway to explore whether either membrane molting or TCR internalization is coupled with the conversion to the memory phenotype.

## MATERIALS AND METHODS

### Antibodies and reagents

Antibodies for Na^+^K^+^ATPase (ab7671), GFP (ab290), TSG101 (ab125011), Rab11 (ab3612), and Annexin V/propidium iodide (PI) stain kit were purchased from Abcam (Cambridge, MA, USA). Antibody for CD3 ζ (sc1239) was purchased from Santa Cruz Biotechnology (Dallas, TX, USA). Antibodies against β-actin (4967L), phospho-Zap70 (2701S), Notch1 (3608S), phospho-PKCδ/θ (9376S), phospho-p44/42 MAPK (Erk1/2, 4376S), phospho-p38 MAPK (9215S), and phospho-Akt (4058S), in addition to anti-rabbit IgG-HRP (7074S) and anti-mouse IgG-HRP (7076S), were purchased from Cell Signaling Technology (Danvers, MA, USA). Anti-TCRβ (H57-597) was purchased from Bio-X-Cell (West Lebanon, NH, USA). Anti-LFA-1 antibody (M17/4) was purchased from BioLegend (San Diego, CA, USA). Fluorescence-conjugated CD3e, CD62L, CXCR4, CD45, CD25, CD11c, CD69, CD4, and CD8 were purchased from eBioscience (San Diego, CA, USA). Flow cytometry submicron particle size reference kit (F13839), Fab preparation kit, CellMask™ Green Plasma Membrane Stain (CMS), CellTrace™ Violet (CTV) cell proliferation kit, CMRA-Orange, Cholera Toxin Subunit B (Recombinant), Pierce™ FITC Antibody Labeling Kit, Alexa Fluor™ 594 or 647 Protein Labeling Kit, and NucSpot Far-Red were purchased from Thermo Fisher Scientific (Waltham, MA, USA). Anti-human CD28 antibody and recombinant mouse ICAM-1/CD54 His-tag were purchased from R&D Systems (Minneapolis, MN, USA). Hybridoma cell lines for mouse anti-CD3 (145-2C11; CRL-1975), mouse anti-CD28 (PV1; HB-12352), and anti-human CD3 (OKT3; CRL-8001) were purchased from the American Type Culture Collection (Manassas, VA, USA). OVA peptide fragments (323–339 or 257–264) were purchased from GeneScript (San Francisco, CA, USA). Reverse transcription PCR premix and restriction enzymes were purchased from Enzynomics (Daejeon, Korea). PLL, MG132, latrunculin A (LatA), cytochalasin D (CytD), chlorpromazine (CPZ), jasplakinolide (JPK), CK636, colchicine (COL), chloroquine (CQ), dynein (DYN), and concanavalin A (ConA) were purchased from Sigma (St. Louis, MO, USA). Vps4a- and Vps4b-targeting and scrambled control siRNAs were purchased as a pool of four si-RNA duplexes from Dharmacon (Lafayette, CO, USA).

### Cells

Jurkat T cells or COS-7 cells expressing Vstm5 fused with GFP (V5G) have been described previously (*13*). Naive CD3^+^, CD4^+^, or CD8^+^ T cells from C57BL/6, *OTII TCR*, or *OTI TCR* C57BL/6 mice, respectably, were purified from mouse spleens and lymph nodes by negative selection using a T-cell enrichment column (R&D Systems). For the generation of OTI or OTII T blast cells, cells from the spleen and lymph nodes of *OTI* or *OTII TCR* mice were stimulated with 2 µg/mL of plate-bound anti-CD3/CD28 antibodies in the presence of rIL-2 (100 U/mL) for 2 days. For the establishment of B-cell blasts, CD19^+^ cells were purified from C57BL/6 wild-type mice using the EasySep magnetic separation system (Stemcell Technologies, Vancouver, Canada) and were activated with lipopolysaccharide (10 μg/mL) for 3 days in complete RPMI 1640 medium. For the isolation of bone marrow– derived dendritic cells (BMDCs), bone marrow was flushed from the femur and tibia bones, and 5 × 10^6^ cells were cultured in 10 mL of RPMI 1640 medium supplemented with 20 ng/mL recombinant murine granulocyte-macrophage colony-stimulating factor (GM-CSF) for 9 days. GM-CSF was added at every 3 days. To generate fixed DCs, 1 × 10^6^ DCs were placed on a fibronectin-coated plate (10 µg/mL) for 2 h, fixed with 0.4% paraformaldehyde for 60 s at room temperature, washed three times with 1× PBS, and stored at 4°C until use.

### Animals

C57BL/6 wild-type, *OTI*, or *OTII* transgenic mice (C57BL/6 background) were purchased from Jackson Laboratory (Bar Harbor, ME, USA). All mice were bred under specific pathogen-free conditions on a 12-h light/dark cycle at the Laboratory Animal Resource Center located in Gwangju Institute of Science and Technology. During the experiments, the mice were euthanized according to the Association for Assessment and Accreditation of Laboratory Animal Care International (AAALAC) guidelines. In general, sex-matched, 8-week-old mice were used.

### T-cell activation

Naive CD3^+^ T cells were activated by incubation on plate-immobilized anti-CD3 (2–10 µg/mL)/CD28 (2 µg/mL) (immobilized conditions, iAb) or by treatment with soluble anti-CD3 (2–10 µg/mL)/CD28 (2 µg/mL) followed by anti-hamster IgG as a secondary antibody (soluble conditions, sAb) for 3 h. Cells were then harvested, washed, and incubated for further 48–72 h. In some experiments, naive CD3^+^ T cells were stimulated with anti-CD3/28 sAb (10 µg/mL/2 µg/mL each), ConA (1 µg/mL), or PMA/ionomycin (200 nM/1 µM) in the presence of coated PLL (1×, 5 × 10^−4^%; 10×, 5 × 10^−3^%), recombinant ICAM-1 (10 µg/mL), anti-CD62L antibody (10 µg/mL), or fixed DCs. For the lipid bilayer, the density of ICAM-1 was 50–500 molecules/μm^2^, unless otherwise noted, and a density of 200 molecules/μm^2^ was used. OTII CD4^+^ or OTI CD8^+^ T cells were stimulated on lipid bilayers presenting 1 μg/mL of OVA_323-339_/I-A^b^ or _257-264_ H-2^K^ with or without ICAM-1. In some experiments, OTII CD4^+^ or OTI CD8^+^ T cells were activated by coincubation with pOVA_323– 339_ or _257–264_-pulsed B cells or DCs.

### Cell transfection and viral infection

For retroviral transduction, mouse CD4^+^ T cells obtained from OTII TCR transgenic mice were incubated in 2 μg/mL anti-CD3/28-coated culture plates with 100 U/mL rIL-2 for 48 h. Retroviral particles were generated by transfection with V5G_pMSCV or TCRζ_GFP_pMSCV along with the pCL-Eco packaging vector using Lipofectamine 2000 (Invitrogen, Carlsbad, CA, USA). After 48 h, viral supernatants were harvested, mixed with 1 × 10^6^ mouse T cells, placed on 20 μg/mL RetroNectin (Clontech, Mountain View, CA, USA)-coated 12-well plates, and spin-infected at 2000 × *g* for 90 min at 25°C with rIL-2 (100 U/mL). The transduced T cells were maintained with fresh mouse T media with rIL-2 and expanded for 3 days. Knockdown of Vps4a/b was performed using Amaxa Nucleofector II and the Mouse T-Cell Nucleofector Kit (Lonza, Basel, Switzerland) according to the manufacturer’s instructions. For transient transfection, COS-7 or Jurkat T cells were transfected with V5G_N1 using Lipofectamine 2000 or Amaxa Nucleofector II, respectively.

### Reverse transcription PCR and real-time quantitative (q) PCR

Total RNA was isolated from cells using the TRI reagent (Molecular Research Center, Cincinnati, OH, USA) and reverse-transcribed using RT-Premix (Intron Biotechnology). PCR was performed with the following primers (the respective forward and reverse pairs are indicated): mouse c-Myc, 5′-TTGAAGGCTGGATTTCCTTTGGGC-3′ and 5′-TCGTCGCAGATGAAATAGGGCTGT-3′; and mouse GAPDH, 5′-GCACAGTCAAGGCCGAGAAT-3′ and 5′-GCCTTCTCCATGGT GGTGAA-3′. The expression levels of c-Myc were evaluated by qPCR. Amplification was performed in a StepOne real-time PCR system (Applied Biosystems, Norwalk, CT, USA) for continuous fluorescence detection in a total volume of 10 μL of cDNA/control and gene-specific primers using SYBR Premix Ex Taq (TaKaRa Bio). The mRNA levels of the target genes were normalized relative to those of *Gapdh* using the following formula: relative mRNA expression = 2^−(ΔCt of target gene − ΔCt of GAPDH)^, where Ct is the threshold cycle value. In each sample, the expression of the analyzed gene was normalized to that of GAPDH and described as the mRNA level relative to GAPDH.

### Flow cytometry analysis

To quantitate total TCRs under various types of stimulation, cells were stained with TCRβ (H57Fab-FITC, Alexa594, or Alexa647) antibody before or after stimulation, and the mean fluorescence intensity (MFI) was measured by flow cytometry. In general, cells were stained with the indicated antibodies for 1 h in 50 µL of FACS buffer (2% FBS in PBS) at 4°C and washed three times with 1× PBS before analysis. For quantification of the percentage (%) of TCR recovery on surface, the MFI of TCR expression in T cells at −3 h (resting condition) was set as 100%. The recovery index was presented as the MFI at each time point divided by the MFI at −3 h. For the analysis of apoptosis, cells were stained with Annexin V and PI according to the manufacturer’s instructions. Data were acquired using a FACSCanto flow cytometer (Becton Dickinson, Franklin Lakes, NJ, USA) and analyzed using the FlowJo software (TreeStar, Inc., Ashland, OR, USA). For the proliferation assay, naive CD3^+^ T cells were stained with CTV and activated as described in the T-cell activation section; cells were then washed, further cultured with fresh media for 72 h in the presence of rIL-2, and analyzed by flow cytometry. In another experiment, cells at 30 h (CD8^+^T) or 48 h (CD4^+^ T) after stimulation were permeabilized using Cytofix/Cytoperm (Invitrogen) and stained with a mixture of anti-Ki-67 antibody and NucSpot Far-Red for 15 min at room temperature in the dark. The samples were washed with PBS and analyzed by flow cytometry. The percentage of proliferative populations was acquired from the gate in a CTV^+^ population.

### Confocal and TIRF microscopy

To investigate the distribution of TCRβ under the different stimulations (iAb and sAb), naive CD3^+^ T cells were stained with TCRβ (H57Fab-Alexa 594) and PMS-Green for 1 h at 4°C and activated with iAb or sAb for 2 h. For the visualization of TCRβ on T cells or DCs during synapse formation, CTV-labeled OTII CD4^+^ T cells were stained with TCRβ (H57Fab-FITC) and incubated with DCs (CMRA-Orange) in the presence or absence of pOVA_323–339_ for 2 h. In some experiments, T cells were pretreated with anti-LFA-1 blocking antibody (M17/4, 10 µg/mL) for 30 min. For intracellular staining, cells were fixed with 4% paraformaldehyde, permeabilized with 0.1% Triton X-100 in PBS, blocked with 1% BSA/PBS for 1 h, rinsed with PBS, and incubated overnight with anti-ezrin antibody at 4°C. Secondary antibodies were added after washing and incubated in the dark for 1 h at room temperature. For the imaging of TMP generations, OTI CD8^+^ T cells were stained with TCRβ (H57Fab-FITC) and stimulated on the lipid bilayer presenting OVA_257–264_/H-2K^b^/with a different dose of ICAM-1. To examine the effects of actin modulators on T-cell activation, OTII CD4^+^ T cells were stimulated on the lipid bilayer presenting OVA_323–339_/I-A^b^/with ICAM-1 in the presence of actin modulators, such as CK636 (100 μM), Lat A (237 nM), and JPK (100 nM) for 2 h. For actin staining, the cells were fixed with 4% paraformaldehyde, permeabilized with 0.1% Triton X-100 in PBS, and incubated with TRITC-phalloidin in PBS for 30 min at room temperature in the dark. All samples were then imaged using a 100× NA 1.40 oil immersion objective lens and an FV1000 laser scanning confocal microscope (Olympus, Japan) or a Laser Scanning Microscopy VIS Laser module, LSM 880 (Carl Zeiss, Germany).

To monitor the TMP generation during T-cell kynapse, V5G^+^-transduced OTII CD4^+^ T blasts were placed on the lipid bilayer presenting OVA_323–339_/I-A^b^/with ICAM-1 and immediately imaged for long time periods (∼2 h) by TIRFM (IX-81; Olympus, Tokyo, Japan) equipped with a solid-state laser (488 nm, 20 mW; Coherent, Santa Clara, CA, USA).

### Electron microscopy

For transmission electron microscopy, isolated TISs were fixed in suspension at 37°C in a 5% CO_2_ incubator and TISs containing the sample (1 mL) were added to 9 mL of fixative (2.2% glutaraldehyde in 100 mM NaPO_4_ [pH 7.4]) for 2 h. Samples were postfixed in 1% osmium tetroxide, stained *en bloc* with 0.5% uranyl acetate in water, dehydrated in a graded ethanol series, embedded, and thinly sectioned. The sections were stained with 2% uranyl acetate in methanol for 20 min, followed by lead citrate for 5 min, and then observed under a Tecnai G2 electron microscope (FEI, Hillsboro, OR, USA) at 120 kV under low-dose conditions. For SEM, cells were fixed with a 2.5% glutaraldehyde solution and osmium tetroxide for 2 h. Samples were then dehydrated in a graded ethanol series for 30 min, dried, prepared by sputter coating with 1–2-nm gold–palladium, and analyzed using a field-emission SEM (Hitachi, Tokyo, Japan).

### SEM analysis of microvilli number and length

All SEM imaging data, including microvilli around one single cell, were analyzed using the ImageJ software and FiloQuant plugin. For quantification of the number and length of microvilli inner area of the cells, the following six major steps were applied: (1) background subtraction, (2) sharpening and finding edges, (3) smoothing and converting to black and white, (4) closing and filling holes, (5) denoizing and segmenting, and (6) filtering and measuring. The number and length of cell-edge filopodia were automatically analyzed using the ImageJ software and FiloQuant plugin. A distribution scatter plot of the numbers of microvilli and TCR clusters was created using the GraphPad Prism 8 software (San Diego, CA, USA).

### TIS isolation and purification

The purification protocol of TISs was modified according to a previously published study (*13*). A total of 20 million cells were resuspended in serum-free RPMI 1640 medium and incubated on anti-CD3/CD28 antibody-coated plates for 2 h at 37°C. Cells were then removed by shaking with cold PBS, and TISs on culture dishes were dissociated by shaking and gentle pipetting (or scrapping) in the presence of cold PBS containing 10 mM EDTA. All supernatants were subjected to two successive centrifugations at 2000 × *g* for 10 min and 2500 × *g* for 10 min to completely eliminate cells and debris. TISs in PBS were reloaded onto a sucrose gradient of 12 different sucrose concentrations from top to bottom (10%–90% sucrose) and centrifuged at 100,000 × *g* for 16 h. Fractions were then carefully collected at 2 mL each from the bottom of the tube. All fractionated samples were diluted with PBS and subjected to centrifugation at 100,000 × *g* for 90 min to pellet the vesicles. Unless otherwise mentioned, 4–6 consecutive fractions of the 12 fractions were used for western blot, SEM, TEM, and LC-MS/MS analyses.

### Quantitation of TISs by flow cytometry

Cells were stained with the Cell Mask green plasma membrane stain for 30 min at 37°C before isolating TISs and harvested as described in the TMP isolation and purification section. TISs were further stained with the Cell Mask green plasma membrane stain for at least 30 min in the dark at room temperature for clear identification. TMP pellets were suspended with 1 mL of 1× PBS and diluted 10-fold with 1× PBS for flow cytometric analysis. Depending on the preparation method and particular cell types, the appropriate dilution for the TMP samples before analysis was typically 1:10–1:1000 relative to the initial sample when running at a slow sample rate (10 μL/min). All experiments in this study were performed using a 3-laser CytoFLEX Flow Cytometer (Beckman Coulter, Brea, CA, USA) and operated using the CytExpert Software v2.4.0.28 (Beckman Coulter). For quantitation, the configuration was modified for VSSC detection. Briefly, the 405/10 VSSC filter was moved to the V450 channel in the wavelength division multiplexer and the detector configuration was modified in the CytExpert Software to assign the VSSC channel within the wavelength division multiplexer. The event-rate setting was set to high before initiating analyses, tightening the pulse window and thus reducing the background for small particle analyses. Finally, the trigger channel was set to VSSC-Height, and the threshold level was manually set as appropriate for small particles. The optimal threshold setting for the CytoFLEX was determined empirically using the Flow Cytometry Submicron Particle Size Reference Kit at their optimal dilution.

### Western blot analysis

Cells or TISs were lysed in ice-cold lysis buffer (50 mM Tris-HCl pH 7.4, 150 mM NaCl, 1% Triton X-100, 1× complete protease/phosphatase inhibitor cocktail) for 1 h on ice. Lysates were centrifuged at 16,000 × *g* for 25 min at 4°C, and the harvested supernatants were mixed with sodium dodecyl sulfate (SDS) sample buffer (100 mM Tris-HCl pH 6.8, 4% SDS, 20% glycerol, bromophenol blue) and then boiled for 5 min. Proteins were separated on 10%–12% SDS polyacrylamide gels by electrophoresis and transferred onto nitrocellulose membranes using a Trans-Blot SD Semi-Dry transfer cell (Bio-Rad, Hercules, CA, USA). Membranes were blocked with 5% skim milk for 1 h, rinsed, and incubated overnight with primary antibodies in Tris-buffered saline (TBS; 50 mM Tris-HCl pH 7.4, 150 mM NaCl) containing 0.1% Tween 20 (TBS-T) and 3% milk. The excess primary antibody was removed by washing the membrane four times in TBS-T before incubation with the peroxidase-labeled secondary antibody (0.1 μg/mL) for 2 h. Bands were visualized using a WEST-ZOL western blot detection kit and exposed to X-ray film.

### Lipidomic analysis

TISs were harvested from 1 × 10^8^ naive CD3^+^ T cells as described in the TMP isolation and purification section, and the samples were lyophilized before lipid extraction. Then, we isolated several lipid species as described previously (*40*). For lipid quantification, we also conducted an SRM-based analysis using product ions obtained in polarity switching mode of the triple-stage quadrupole MS system. For the quantitative normalization of lipid species between T cells and TISs, the MS peak area of individual lipid species was calculated by dividing the peak area of the internal standard.

### Metabolic studies

Naive CD3^+^ T cells were placed on PLL-coated Seahorse Bioanalyzer XFp culture plates (3 × 10^5^ cells/well) with Seahorse XF RPMI Assay Media (RPMI Medium, pH 7.4, 103576-100; Agilent Technologies, Santa Clara, CA, USA), supplemented with 10 mM glucose (103577-100; Agilent), 1 mM pyruvate (103578-100; Agilent), and 2 mM L-glutamine (103579-100; Agilent) for 1 h at 37°C. Basal rates were taken for 24 min, and 20 μL of dynabead mouse CD3/CD28 T-cell activator (2 × 10^6^ beads/mL; dAb, to mimic immobilized conditions) or sAb (10 μg/mL) was injected into the culture plate and stimulated rates were taken for 2 h. To analyze the metabolic changes under different conditions, naive CD3^+^ T cells were stimulated with iAb, sAb, and sAb on ICAM-1-, PLL-, or CD62L-coated plates for 3 h. In some experiments, the cells were pretreated with various inhibitors (LatA, 237 nM; Cyt D, 200 nM; JPK, 100 nM; COL, 10 μM) for 1 h, washed, and stimulated with sAb for 3 h. Cells were then washed, cultured for another 21 h, and placed on PLL-coated Seahorse Bioanalyzer XFp culture plates with Seahorse XF RPMI Assay Media for 1 h at 37°C before measurement.

### Proteomic analysis by LC-MS/MS

TISs were harvested as described in the TMP isolation and purification section, and the same amount of proteins from the total lysate or TISs was subjected to SDS-PAGE on a 12% polyacrylamide gel. The gel was stained with Coomassie Brilliant Blue R-250 and fractionated according to molecular weight. Tryptic in-gel digestion was conducted according to a previously described procedure (*41*). The digested peptides were extracted with an extraction solution consisting of 50 mM ammonium bicarbonate, 50% acetonitrile, and 5% trifluoroacetic acid (TFA) and then dried. For LC-MS/MS analysis, the samples were dissolved in 0.5% TFA. Tryptic peptide samples (5 μL) were separated using an Ultimate 3000 UPLC system (Dionex, Sunnyvale, CA, USA) connected to a Q Exactive Plus mass spectrometer (Thermo Scientific, Waltham, MA, USA) equipped with a nanoelectrospray ion source (Dionex). Peptides were eluted from the column and directed on a 15 cm × 75 μm i.d. Acclaim PepMap RSLC C18 reversed-phase column (Thermo Scientific) at a flow rate of 300 nL/min. Peptides were eluted by a gradient of 0%–65% acetonitrile in 0.1% formic acid for 180 min. All MS and MS/MS spectra obtained using the Q Exactive Plus Orbitrap mass spectrometer were acquired in the data-dependent top10 mode, with automatic switching between full-scan MS and MS/MS acquisition. Survey full-scan MS spectra (m/z 150–2000) were acquired in the orbitrap at a resolution of 70,000 (m/z 200) after the accumulation of ions to a 1 × 10^6^ target value based on predictive automatic gain control from the previous full scan. The MS/MS spectra were searched with MASCOT v2.4 (Matrix Science, Inc., Boston, MA, USA) using the UniProt human database for protein identification. The MS/MS search parameters were set as follows: carbamidomethylation of cysteines, oxidation of methionines, two missed trypsin cleavages, mass tolerance for a parent ion and fragment ion within 10 ppm, and a p value of <0.01 of the significant thresholds. The exponentially modified protein abundance index (emPAI) was generated using MASCOT, and mol% was calculated according to the emPAI values (*42*). The MS/MS analysis was performed at least three times for each sample.

### RNA isolation and Affymetrix GeneChip microarray

For the preparation of RNA samples, 5 × 10^6^ naive CD3^+^ T cells were stimulated with Lat A (237 nM), sAb, iAb, or sAb+LatA for 3 h, washed, and harvested. For the 24-h sample, the cells were washed and further cultured for 21 h after stimulation. In some analysis, cells were stimulated with ConA, PLL, or ConA+ PLL. Total RNA was isolated using an RNeasy Plus Mini Kit (Qiagen, Hilden, NRW, Germany). cDNA was synthesized using the GeneChip Whole Transcript amplification kit purchased from Thermo Fisher Scientific (Waltham, MA, USA). cDNA was fragmented using terminal deoxynucleotidyl transferase and labeled with biotin. The labeled target DNA was hybridized for 16 h at 45°C. Finally, the hybridization signal was calculated using the Affymetrix^®^ GeneChip™ Command Console Software. Gene enrichment and clustering were performed using KEGG (www.genome.jp/kegg/), and a functional analysis was subjected to Gene Ontology (www.geneontology.org/).

### Expression profiling of microarray and pathway analysis

Raw data of the microarray were normalized using the robust multi-average method (*43*), and PCA was performed on the normalized expression profiles of all samples using the R prcomp package. Using the fold change values, we performed rank-based pathway enrichment tests from the R gProfiler (Gene Ontology-based). For in-depth metabolic pathway enrichment tests, we extracted genes belonging to metabolic pathways (i.e., metabolic subsystems) from high-quality genome-scale metabolic models of mouse (MMR version 1) (*26, 27*) and performed pathway enrichment tests using the R fgsea package.

### Statistics

Student’s *t*-test and one-way analysis of variance, corrected for all pairwise comparisons, were performed using the GraphPad Prism software version 8.1.2 (GraphPad, San Diego, CA) between two and more different groups. A p value of <0.01 was considered to be statistically significant. Data are presented as mean ± SD or standard error of the mean.

## Author contributions

J-S. P. conceived the study; J-S. P., J-H.K., W-C.S., and K-S.L. designed and performed the experiments; S.L. and C-H.K. analyzed the data; H.R.K. and C.D.J. wrote and finalized the manuscript.

## Data availability

All data needed to evaluate the conclusions in the paper are present in the paper and/or the Supplementary Materials.

## SUPPLEMENTARY MATERIALS

**Fig. S1.**
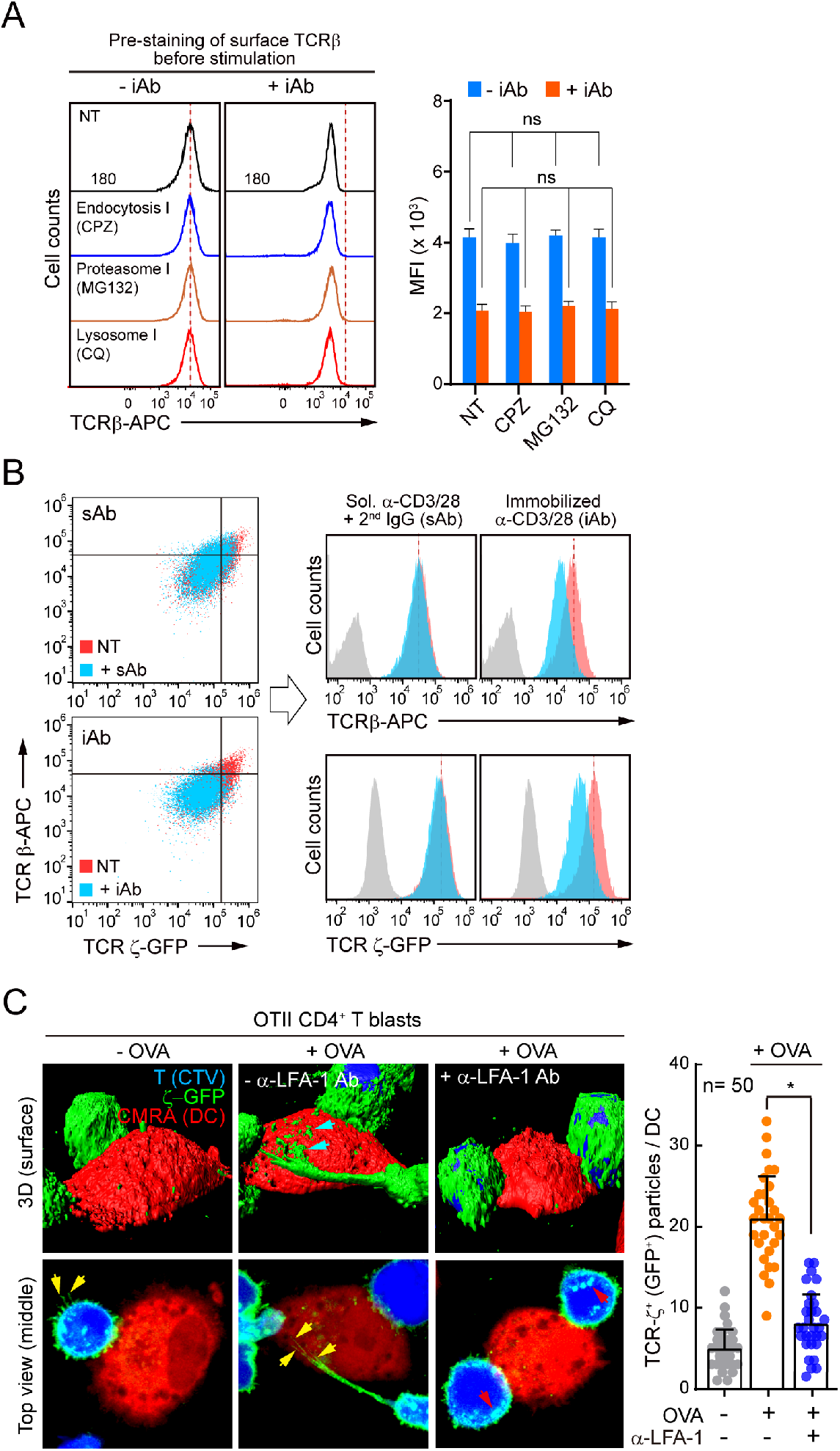
T-cell activation on anti-CD3/CD28 iAb triggers TCR release. (A) Mouse CD3^+^ T cells were stained with anti-TCRβ and treated with various inhibitors, such as chlorpromazine (CPZ, 20 μM), MG132 (20 μM), or chloroquine (CQ, 20 μM), and activated on plate-immobilized (iAb) anti-CD3/CD28 antibodies for 3 h (continued from Fig. 1b). The mean fluorescence intensity (MFI) was measured by flow cytometry. (B) OTII CD4^+^ T cells expressing TCRζ_GFP were stained with anti-TCRβ (H57Fab-Alexa647) and activated with iAb or soluble anti-CD3/CD28 antibodies (sAb) for 3 h. The MFI of each fluorescence was determined by flow cytometry. Results are representative of three independent experiments. (C) CTV-labeled OTII CD4^+^ T cells expressing TCRζ_GFP were incubated with 1 μg/mL OVA_323–339_-pulsed DCs (CMRA-Orange) in the absence or presence of anti-LFA-1 blocking antibody (10 μg/mL). Cyan arrowheads indicate TCRζ_GFP^+^ particles separated from OTII CD4^+^ T blasts. Yellow and red arrowheads indicate the TCRζ_GFP signals located in the microvillar projections or inside cells via internalization. TCRζ_GFP^+^ particles per single DC were quantitated using Imaris. Data represent the mean ± SEM of three independent experiments. ns, not significant, \**p* < 0.05.

**Fig. S2.**
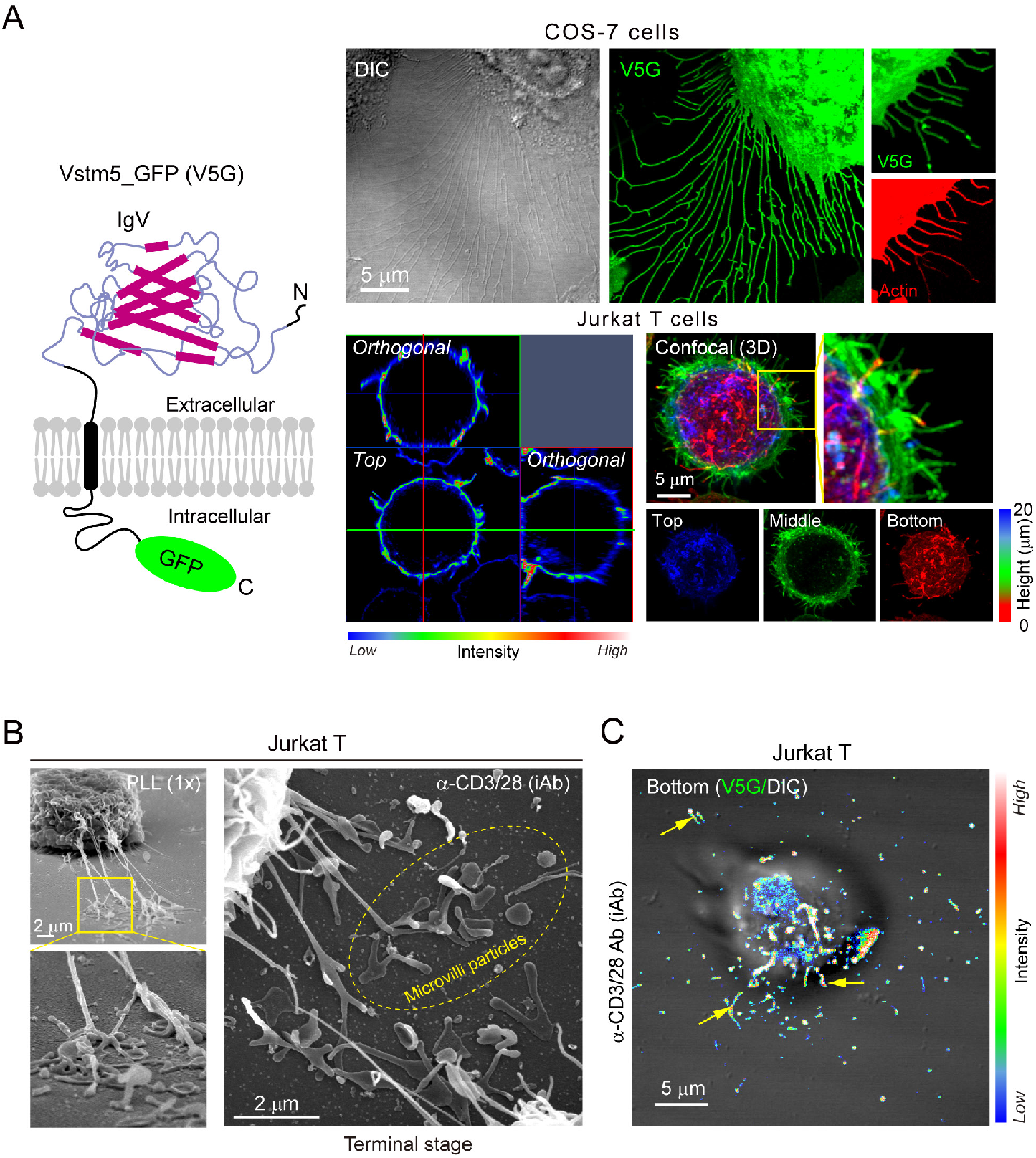
V5G is located at the finger-like membrane-protrusive region: microvilli. (A) Schematic representation of Vstm5 fused with GFP (V5G) structure on the cell membrane and subcellular localization of V5G in COS-7 or Jurkat T cells. V5G-expressing COS-7 cells were fixed and stained with phalloidin-TRITC (top). V5G in Jurkat T cells was represented as pseudo-color coding according to the fluorescence intensity (bottom). (B) SEM evidence of TMP release from Jurkat T cells. Jurkat T cells were incubated on PLL or stimulated with iAb for 60 min (terminal stage). (C) V5G^+^ Jurkat cells were stimulated with iAb for 3 h and observed by confocal microscopy. V5G in microvilli was represented with pseudo-color coding according to the fluorescence intensity. Yellow arrowheads indicate the separated large TMPs.

**Fig. S3.**
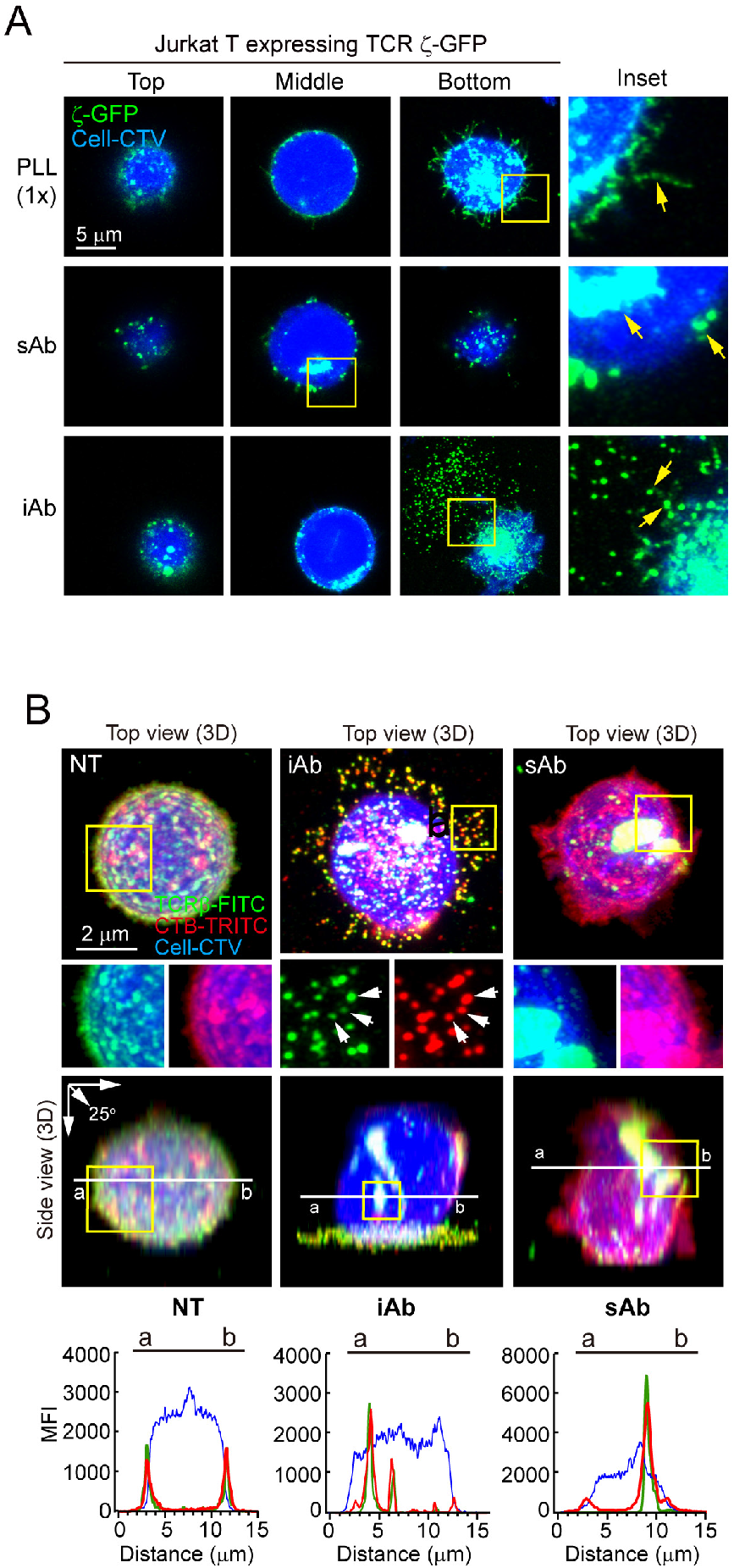
TCR ζ^+^ and CTB^+^ nanoscale particles are released from both human and mouse T cells. (A) Jurkat T cells expressing TCRζ_GFP were stimulated with anti-human anti-CD3/CD28 iAb or sAb and observed by confocal microscopy. Yellow arrowheads indicate the internalized TCRζ_GFP or separated TCRζ_GFP^+^ particles. (B) Colocalization of TCRβ^+^ (FITC) and CTB^+^ (TRITC) signals in released particles from mouse CD4^+^ T cells activated for 3 h on anti-CD3/CD28 iAb or sAb.

**Fig. S4.**
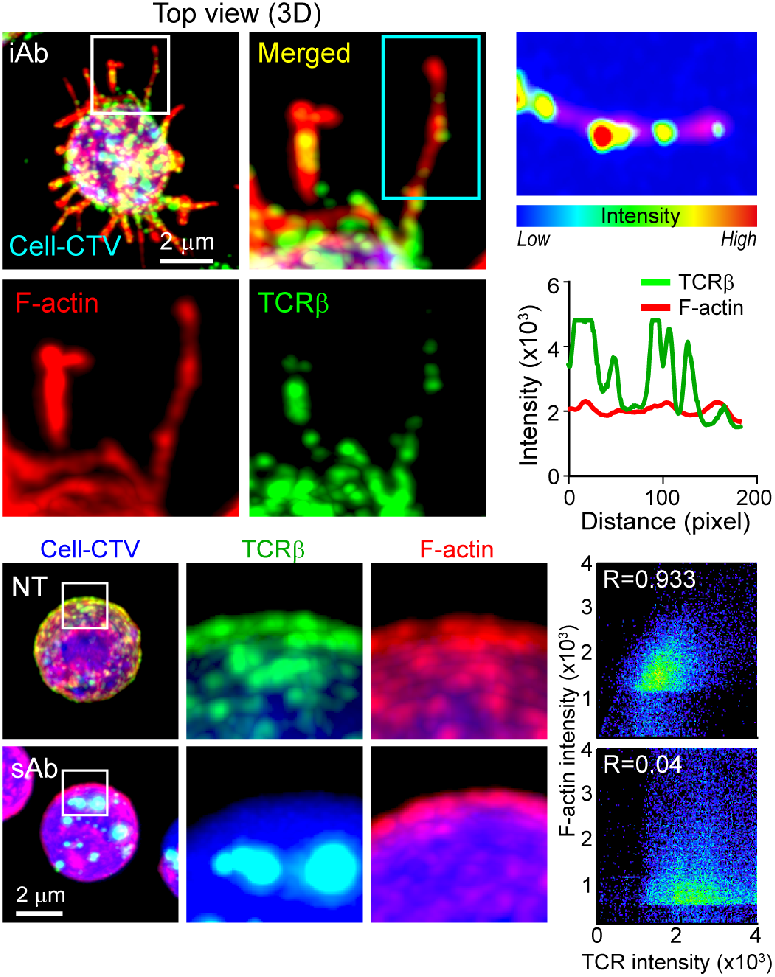
TMPs released via F-actin-enriched microvilli. Naive CD3+ T cells stained with anti-TCRβ (H57Fab-Alexa594) were stimulated (continued from Fig. 3B), fixed, permeabilized, and stained with phalloidin-TRITC. Localization of TCRβ and F-actin was analyzed by confocal microcopy.

**Fig. S5.**
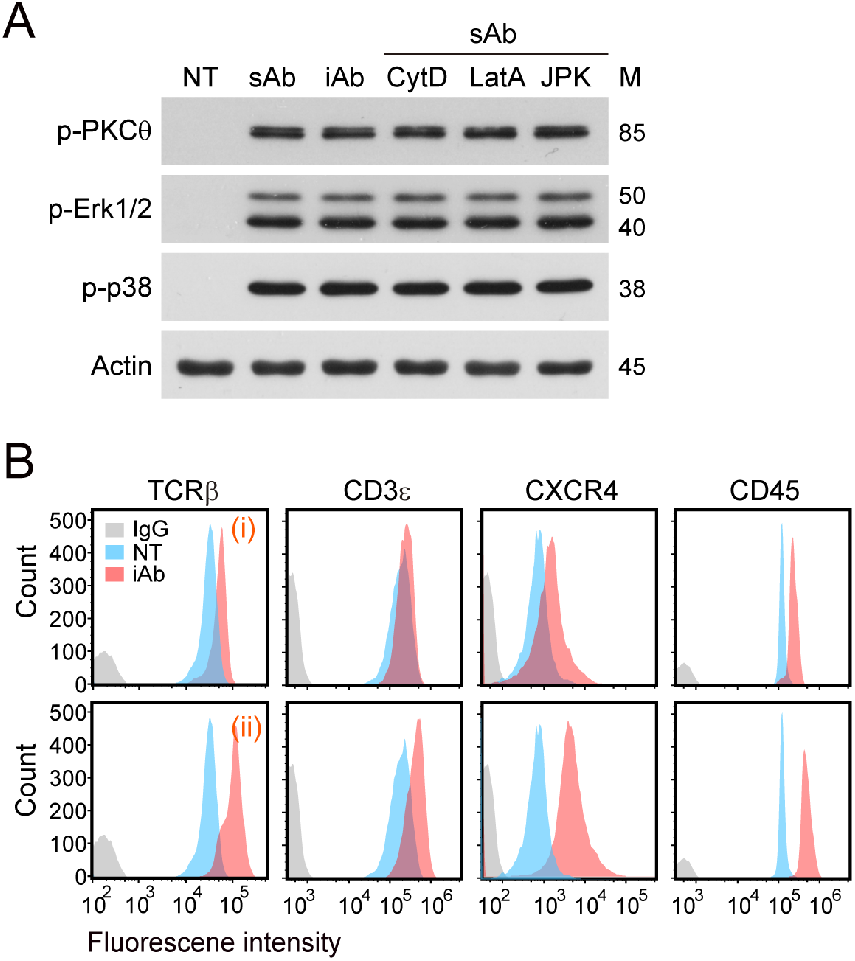
Trogocytic molting of T-cell microvilli increases cell size and surface protein expression. (A) Naive CD3^+^ T cells were activated by anti-CD3/CD28 iAb or sAb for 3 h in the presence or absence of actin-modulating drugs, and the TCR distal signaling pathway was analyzed by western blotting. (B) Cells from Fig. 4B gating (i) and (ii) were analyzed for levels of surface proteins potentially enriched or excluded in microvilli. Results are representative of three independent experiments.

**Fig. S6.**
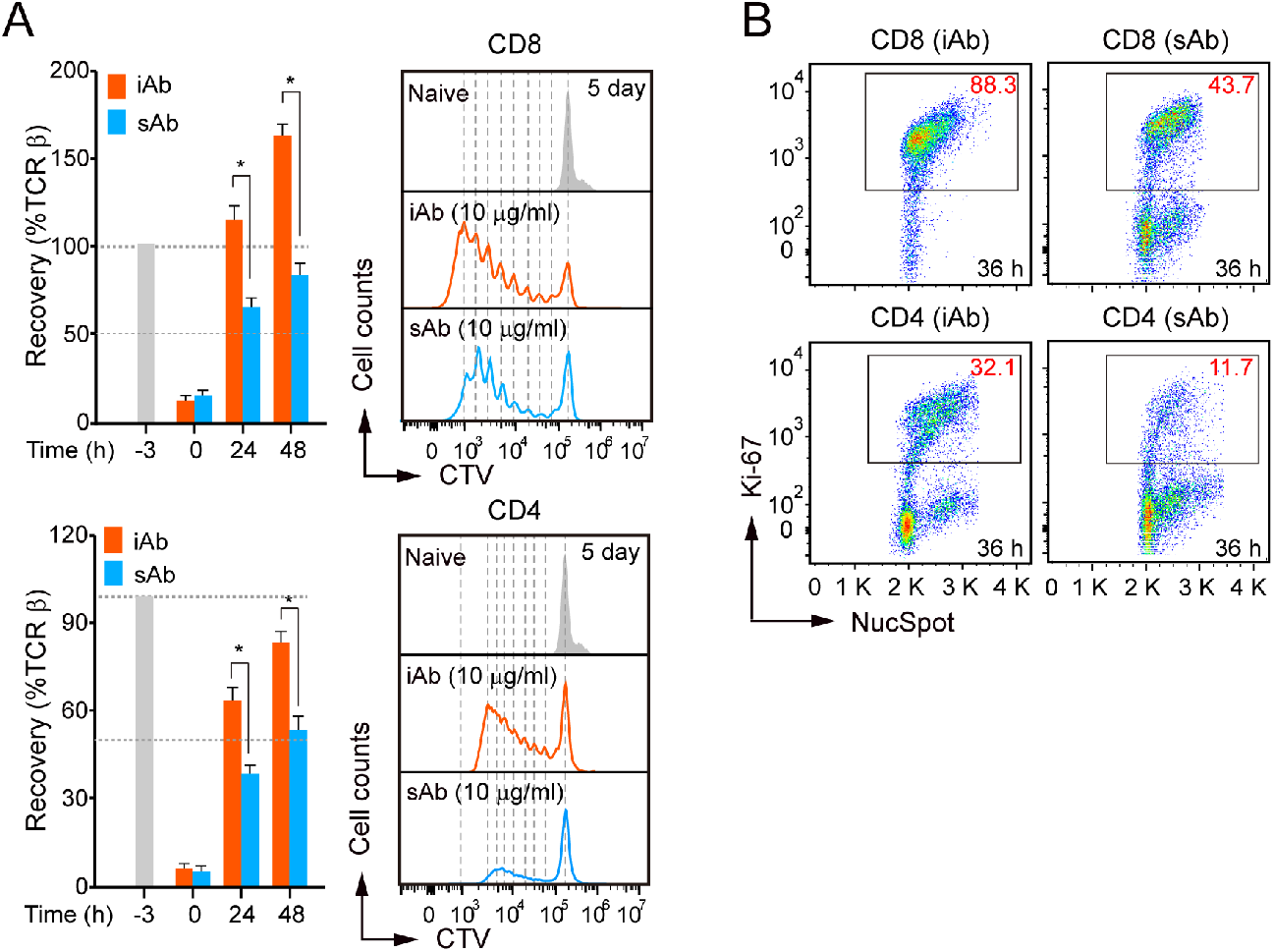
Trogocytic molting of microvilli enhances TCR recovery and T-cell proliferation in CD4^+^ and CD8^+^ T cells. (A and B) CTV-labeled naive CD4^+^ or CD8^+^ T cells were stimulated with anti-CD3/CD28 iAb or sAb for 3 h, washed, and further incubated for the indicated time points. TCR recovery on the cell surface and cell division (A) and the proliferating populations were determined by Ki-67/NucSpot-double-positive staining at 36 h after stimulation (B). Results are representative of three independent experiments. \**p* < 0.05.

**Fig. S7.**
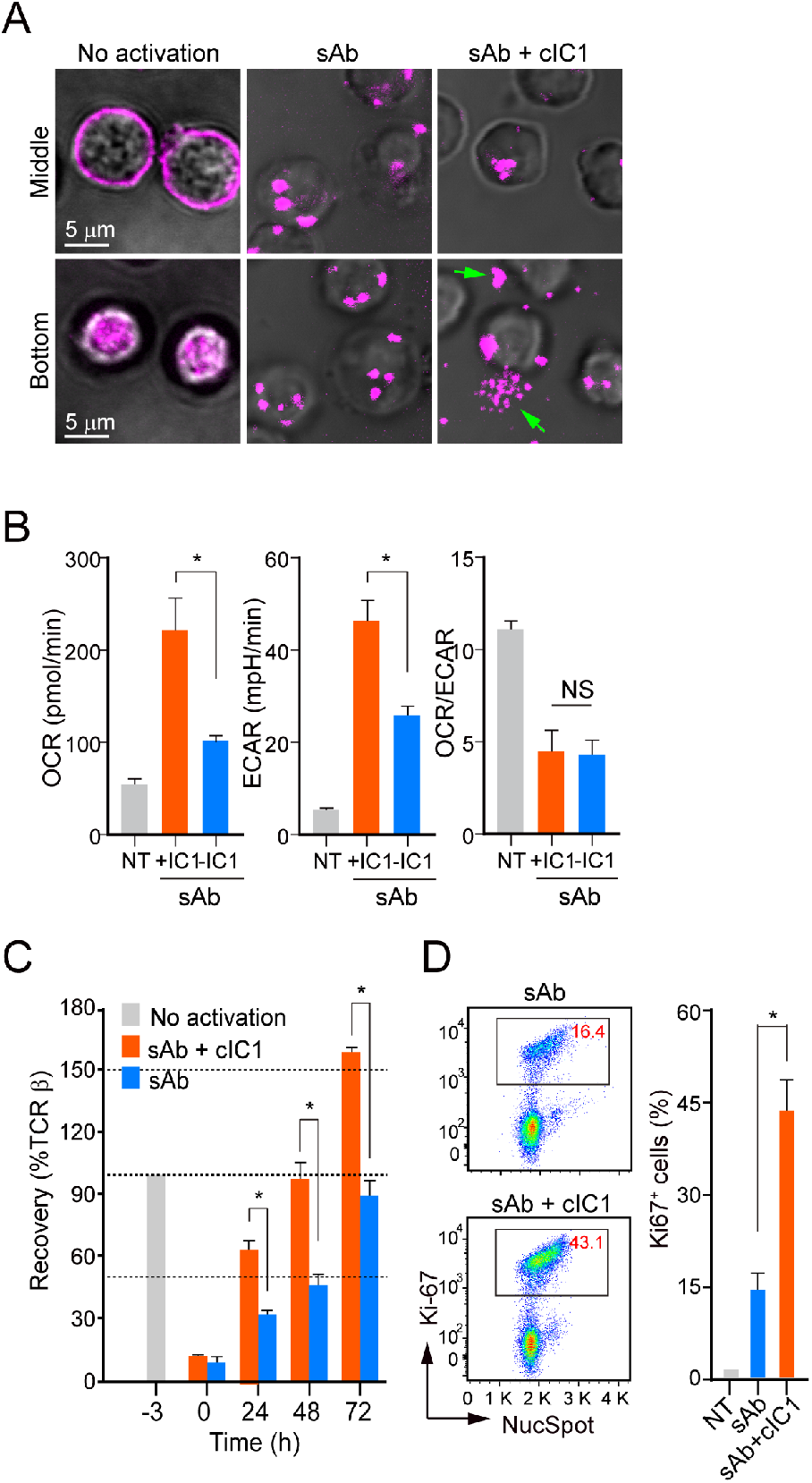
Release of TCR^+^ particles requires both TCR activation and adhesion strength. (A) CD3^+^ T cells were stained with anti-TCRβ-Fab-Alexa 594 at 4°C and stimulated for 3 h with sAb in the presence or absence of coated ICAM-1 (cIC1, 10 μg/mL). Green arrowheads indicate released TCRβ^+^ particles (objective 100×). (B) OCR and ECAR of CD3^+^ T cells at 48 h. \**p* < 0.01. (C) The surface recovery of TCRs. \**p* < 0.01. (D) Ki-67/NucSpot-double-positive cells at 48 h of post stimulation. NT, non-treated; cIC1, coated ICAM-1; NS, not significant. \**p* < 0.01. Values are mean ± SE.

**Fig. S8.**
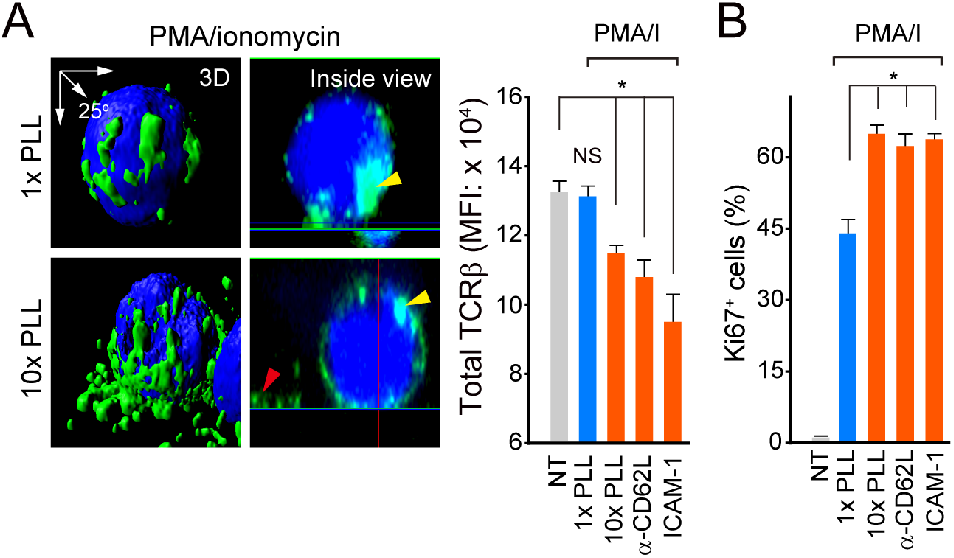
Microvilli release enhances T-cell proliferation by PMA/ionomycin. CTV-labeled naive CD3^+^ T cells were stained with anti-TCRβ (H57Fab-FITC) and stimulated with PMA/ionomycin (200 nM/1 µM) in the presence of indicated conditions for 3 h (continued from Fig. 5H). (A) Distribution of TCRβ^+^ signals (GFP) and the MFI of TCRβ on the T-cell surface were analyzed. (B) The proliferating cells were determined at 48 h after stimulation. Internalized (yellow) and separated (red) TCRβ signals were indicated. Data represent the mean ± SEM of three independent experiments. ns, not significant, \**p* < 0.05.

**Fig. S9.**
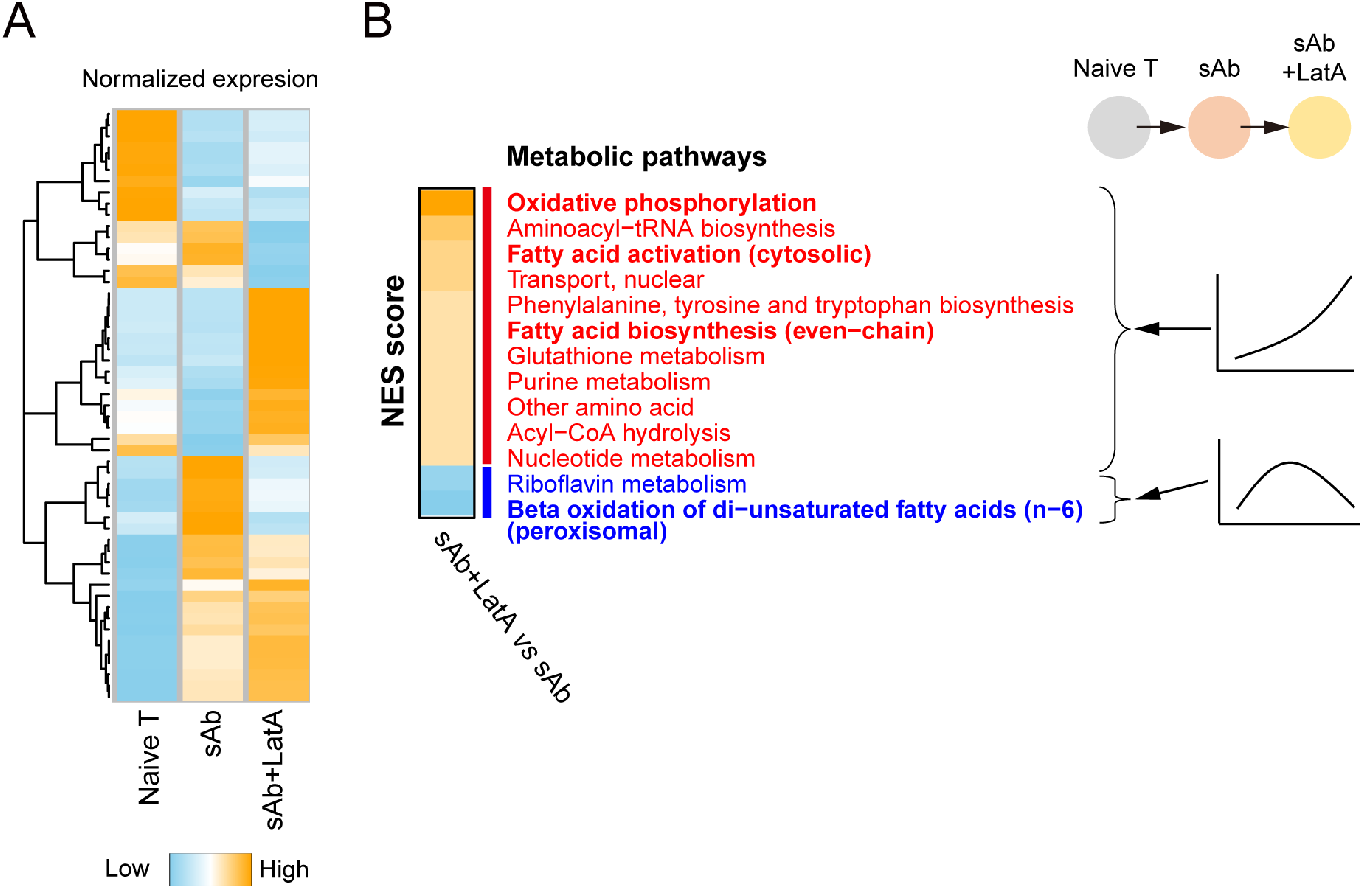
Enrichment tests of metabolic pathways of T cells. (A) Normalized enrichment scores (NES) of metabolic pathways (R fgsea package) were calculated from the fold changes between sAb+LatA-activated T cells and sAb-activated T cells (at 3 h). (B) Increased expression of metabolic pathways, including OXPHOS, FAS, and nucleotide metabolism, and decreased expressions of metabolic pathways, including FAO, were observed. Pathways of the heatmap were selected based on enrichment test p values (<0.05).

**Fig. S10.**
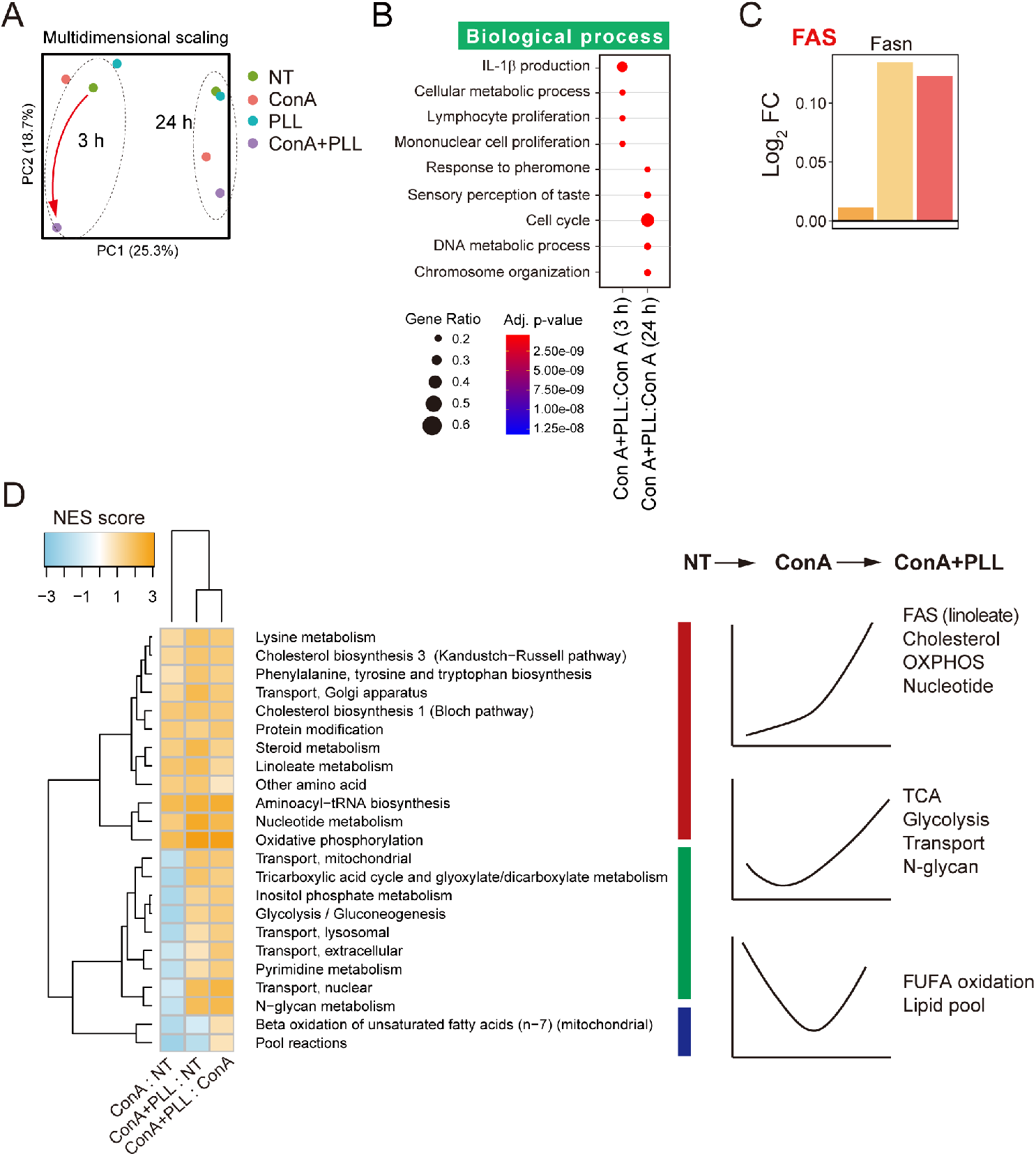
Transcriptome analysis of naive T cells and ConA- and ConA plus PLL-activated naive T cells. (A) Principal component analysis (PCA) of transcriptome. Trajectory was changed from naive T cell to PLL-, ConA-, and ConA+PLL-activated (3 and 24 h) T cells. The changes were time-dependent, observing different clusters by time. (B) Enriched biological pathways (Gene Ontology) of differentially expressed genes. (C) Bar graphs show the fold changes (log2) of representative genes in FAS pathways. (D) Normalized enrichment scores (NES) of metabolic pathways (R fgsea package) were calculated from the fold changes among ConA vs. naive T, ConA+PLL vs. naive T, and ConA+PLL vs. ConA.

**Video S1. T cells do not release TCRβ^+^ fluorescence signals in the absence of antigen on DCs.** OTII CD4^+^ T cells were stained with anti-TCRβ (H57Fab-FITC) and CTV (cytosol), and the cells were incubated for 3 h with DCs (CMRA-Orange) without antigen peptide. Images were 3D-reconstituted, and the IsoSurface module of Imaris was applied. *Abbreviations*: CTV, CellTrace violet; DC, dendritic cells; OVA, ovalbumin; 3D, three-dimensional. 27.3 MB

**Video S2. DCs acquire TCRβ^+^ fluorescence signals from T cells in an antigen- and adhesion-dependent manner.** OTII CD4^+^ T cells were stained with anti-TCRβ (H57Fab-FITC) and CTV (cytosol), and the cells were incubated with 1 μg/mL OVA_323–339_-pulsed DCs (CMRA-Orange) for 3 h. Images were 3D-reconstituted, and the IsoSurface module of Imaris was applied. *Abbreviations*: CTV, CellTrace violet; DC, dendritic cells; OVA, ovalbumin; 3D, three-dimensional. 35.6 MB

**Video S3. Anti-LFA-1 antibody blocks the release of TCRβ^+^ particles from activated T cells on DCs.** OTII CD4^+^ T cells were prestained with anti-TCRβ Fab-FITC and CTV (cytosol), and then the cells were incubated with 1 μg/mL OVA_323–339_-pulsed DCs (CMRA-Orange) in the presence of anti-LFA-1 blocking antibody (10 μg/mL). Images were 3D-reconstituted, and the IsoSurface module of Imaris was applied. *Abbreviations*: CTV, CellTrace violet; DC, dendritic cell; LFA, leukocyte function-associated antigen; OVA, ovalbumin. 35.5 MB

**Video S4. Release of V5G^+^ microvilli particles from OTII CD4^+^ T cells during migration on the lipid bilayers presenting OVA_323–339_/I-A^b^/ICAM-1.** *Abbreviations*: ICAM, intercellular adhesion molecule; V5G, Vstm5 fused with green fluorescent protein. 8.3 MB

**Video S5. Colocalization of TCRβ^+^ (FITC, green) and CTB^+^ (TRITC, orange) signals on the surface of unstimulated naive T cells.** *Abbreviations*: CTB, cholera toxin B subunit; V5G, Vstm5 fused with green fluorescent protein. 14.1 MB

**Video S6. Colocalization of TCRβ^+^ (FITC, green) and CTB^+^ (TRITC, orange) signals in the cytosol of sAb-stimulated T cells.** *Abbreviations*: CTB, cholera toxin B subunit; sAb, soluble anti-CD3/CD28 antibody. 14.0 MB

**Video S7. Colocalization of TCRβ^+^ (FITC, green) and CTB^+^ (TRITC, orange) signals in the released particles or inside the cytosol of iAb-stimulated CD4^+^ T cells.** *Abbreviations*: CTB, cholera toxin B subunit; iAb, immobile anti-CD3/CD28 antibody. 14.1 MB.

